# Equivalent excitability through different sodium channels and implications for the analgesic efficacy of selective drugs

**DOI:** 10.1101/2022.10.04.510784

**Authors:** Yu-Feng Xie, Jane Yang, Stéphanie Ratté, Steven A. Prescott

## Abstract

Nociceptive sensory neurons convey pain-related signals to the CNS using action potentials. Loss-of-function mutations in the voltage-gated sodium channel Na_V_1.7 cause insensitivity to pain (presumably by reducing nociceptor excitability) but efforts to treat pain by inhibiting Na_V_1.7 pharmacologically have largely failed. This may reflect the variable contribution of Na_V_1.7 to nociceptor excitability. Contrary to claims that Na_V_1.7 is necessary for nociceptors to initiate action potentials, we show that nociceptors can achieve equivalent excitability using different combinations of Na_V_1.3, Na_V_1.7, and Na_V_1.8. Selectively blocking one of those Na_V_ subtypes reduces nociceptor excitability only if the other two subtypes are weakly expressed. For example, excitability relies on Na_V_1.8 in acutely dissociated nociceptors but responsibility shifts to Na_V_1.7 and Na_V_1.3 by the fourth day in culture. A similar shift in Na_V_ dependence occurs in vivo after inflammation, impacting ability of the Na_V_1.7-selective inhibitor PF-05089771 to reduce pain in behavioral tests. Flexible use of different Na_V_ subtypes exemplifies degeneracy – equivalent function using different components – and compromises the reliable modulation of nociceptor excitability by subtype-selective inhibitors. Identifying the dominant Na_V_ subtype to predict drug efficacy is not trivial. Degeneracy at the cellular level must be considered when choosing drug targets at the molecular level.

**SIGNIFICANCE STATEMENT:** Nociceptors can achieve equivalent excitability using different sodium channel subtypes. The analgesic efficacy of subtype-selective drugs hinges on which subtype controls excitability. This contingency likely contributes to poor clinical outcomes.

## INTRODUCTION

Chronic pain affects between 11 and 40% of the population worldwide (1). Neuropathic pain, which is pain arising from damage to the somatosensory nervous system, is particularly hard to treat with only 30% of patients achieving moderate (≥30%) relief using available treatments (2, 3). New treatments are needed but a meagre 11% of analgesic drugs entering phase 1 trials are ultimately approved (4), triggering debate about why basic science discoveries are not yielding improved clinical outcomes (5). Suggested explanations include flaws in preclinical animal testing (6, 7) or clinical trial design (8) but biological explanations must also be considered. For example, degeneracy – the ability of a biological system to achieve equivalent function using different components (9) – complicates modulation of neuronal excitability by allowing changes in diverse ion channels to potentially subvert the therapeutic effect of a drug targeting a particular channel (10). These explanations are not mutually exclusive but degeneracy continues to receive little consideration.

Like most neurons, nociceptive sensory neurons (nociceptors) rely on spikes to transmit information. Their excitability is thus critical for relaying information to the CNS. Nociceptor excitability is increased in many pathological pain conditions and the resultant increase in afferent input drives chronic pain (11-13). Neuronal excitability depends on the complex interplay between diverse ion channels (14-16) but some channels seem to be particularly important for pain. For instance, loss- or gain-of-function mutations in the gene *SCN9A*, which encodes the voltage-gated sodium channel Na_V_1.7, cause congenital insensitivity to pain (CIP) or painful neuropathies, respectively (17-19); for review see (20). In rodents, nociceptor-specific deletion of Na_V_1.7 abolishes acute and inflammatory pain (21) but not neuropathic pain (22, 23). Neuropathic pain is blocked by deleting Na_V_1.7 globally, including from sympathetic neurons (24, 25), though not if the deletion is induced in adulthood (26). Furthermore, loss-of-function mutations in Na_V_1.7 do not consistently reduce nociceptor excitability (see Discussion) and the associated insensitivity to pain involves increased opioid signaling (27, 28), consistent with naloxone’s ability to restore pain sensitivity in CIP patients (27, 29). These observations cast doubt on whether Na_V_1.7 mutations produce CIP by reducing nociceptor excitability, pointing instead to a less direct mechanism that may be harder to reproduce pharmacologically.

Notwithstanding such reservations, several Na_V_1.7-selective drugs have been developed (30-32) but none have yet passed phase 2 clinical trials (33-36). This has been attributed to poor target engagement (35, 37-39) yet prevention of the flare response by PF-05198007, a Na_V_1.7-selective inhibitor, argues that at least some Na_V_1.7 channels are blocked (40). But CIP patients exhibit a normal flare response (41), suggesting that their C fibers compensate for chronic loss of Na_V_1.7 channels. Other Na_V_1.7-selective inhibitors have struggled in phase 1 trials because of autonomic side effects (e.g. (42)), as might be expected if those drugs block Na_V_1.7 channels on sympathetic neurons, which is apparently necessary to prevent/reverse neuropathic pain (see above). But CIP patients exhibit normal autonomic function (17, 41), suggesting that their sympathetic neurons also compensate for chronic loss of Na_V_1.7 channels. In those patients, might similar compensation occur in nociceptors and restore pain, only for that effect to be masked by enhanced opioid signaling (see above)? Descriptions of Na_V_1.7 as “the” threshold channel imply that it is irreplaceable for nociceptor excitability, consistent on the surface with pain insensitivity due to loss-of-function mutations in Na_V_1.7 but inconsistent with some past electrophysiological data (43, 44). Clarifying whether nociceptors rely on Na_V_1.7 is an unresolved issue important for predicting the analgesic efficacy of Na_V_1.7-selective inhibitors.

A serendipitous observation prompted us to reassess the role of Na_V_1.7 in nociceptor excitability and the implications for drug efficacy. Specifically, we observed that tetrodotoxin (TTX), which inhibits Na_V_1.7 and several other TTX-sensitive (TTX-S) sodium channels, had variable effects in nociceptors, dramatically reducing their excitability in some conditions but not in others. This variability reveals that nociceptors can achieve equivalent excitability using different sodium channel subtypes, some of which are TTX-resistant (TTX-R). We demonstrate that a Na_V_1.7-selective inhibitor produces analgesia only when nociceptor excitability relies on Na_V_1.7. Insofar as increasingly selective drugs are more likely to have their efficacy subverted by degeneracy, our results have profound yet underappreciated implications for target selection and drug development.

## RESULTS

### Equivalent excitability can arise from different voltage-gated sodium (Na_V_) channel subtypes

Small dorsal root ganglion (DRG) neurons (soma diameter <25 µm) tend to spike repetitively when depolarized by current injection (45). In our sample, most small neurons genetically identified as nociceptors (see Methods) spiked repetitively when tested 2-8 hours after dissociation (DIV0) or after 4-7 days in culture (DIV4-7), though the proportion of repetitively spiking neurons increased slightly over that interval (*χ*^2^=4.51, p=0.034, chi-square test) (**Fig. 1A**). Strikingly, 100 nM TTX had no effect on the spiking pattern at DIV0 but converted all but one neuron to transient spiking at DIV4-7. Amongst neurons that spiked repetitively at baseline, TTX reduced the firing rate and increased rheobase only at DIV4-7 (**Fig. 1B**). TTX reduced spike height at DIV0 and DIV4-7, but more so at DIV4-7. There was a significant increase in capacitance and leak conductance density between DIV0 and DIV4-7, but no change in resting membrane potential (**Fig. 1C**). Normalizing leak conductance by capacitance (which increases over time because of neurite growth) disambiguates whether changes in input resistance reflect changes in cell size or membrane leakiness. Consistent with current clamp data, voltage clamp recordings showed that only a small fraction of sodium current is TTX-S at DIV0, whereas nearly all sodium current was blocked by TTX at DIV4-7 (**Fig. 1D**). Previous studies suggested that TTX-R channels play an important role in nociceptor excitability (46-48). Our initial results confirm this for DIV0 but show that their contribution diminishes after a few days in culture, with TTX-S channels becoming dominant by DIV4. Despite this reconfiguring of Na_V_ channels, excitability was remarkably stable, consistent with previous work showing little change in excitability after axotomy despite large (but evidently counterbalanced) changes in TTX-R and TTX-S currents (43, 49). We show later that similar changes develop in vivo following inflammation with consequences for drug efficacy assessed behaviorally (see Fig. 8), suggesting the Na_V_ channel reconfiguration described above is not a trivial epiphenomenon of culturing.

**Figure 1.**
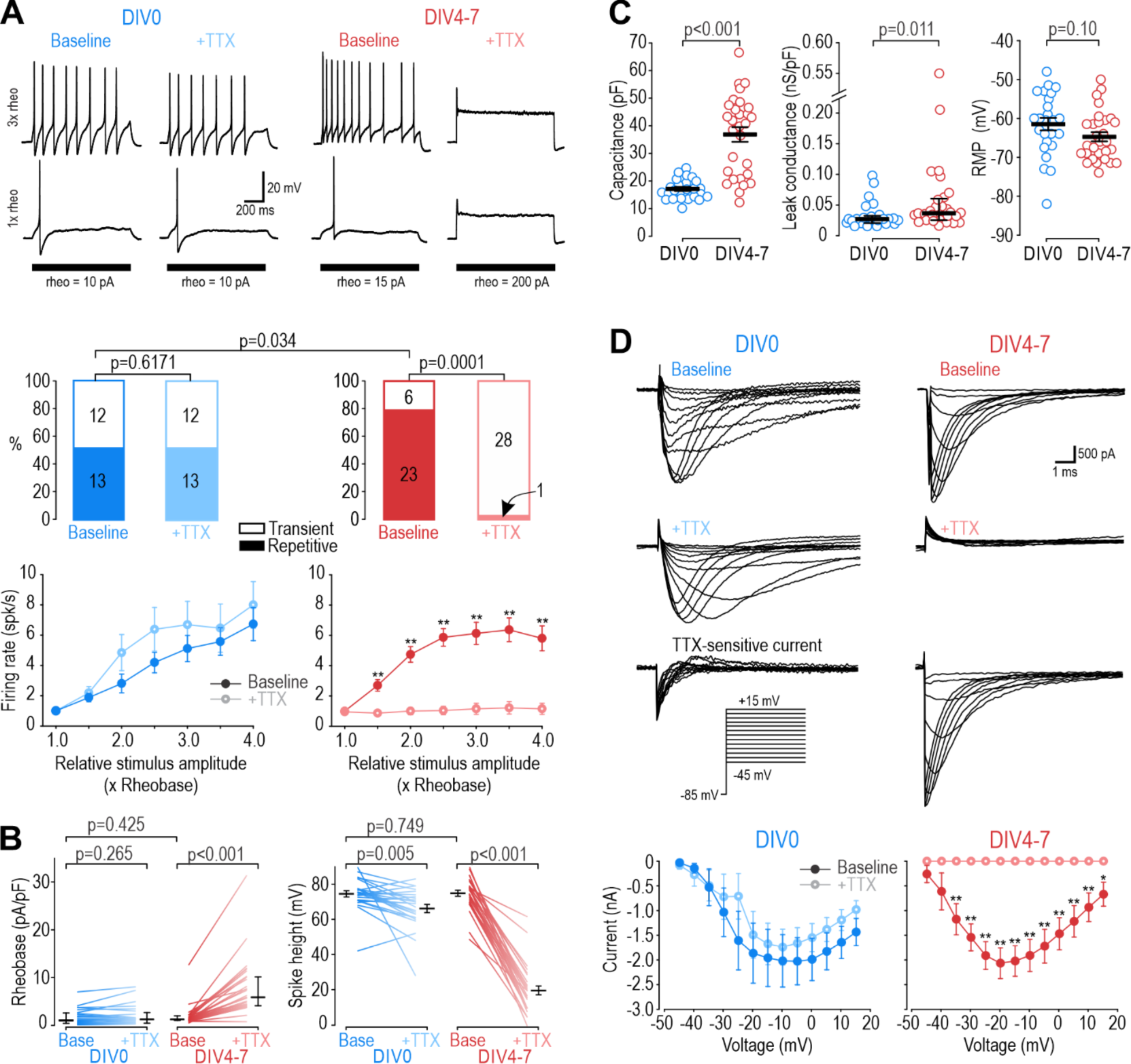
Different Na_V_ subtypes produce equivalent excitability at different days in vitro (DIV). **(A)** Representative responses of small DRG neurons to current injection at rheobase and 3x rheobase when tested on DIV0 (blue) or DIV4-7 (red) before (dark) and after (pale) bath application of 100 nM TTX. At DIV0, TTX did not alter spiking pattern (*χ*^2^=0.25, p=0.617, McNemar test) or significantly reduce firing rate (F_1,72_=1.527, p=0.24, two-way repeated measure (RM) ANOVA; n=13). At DIV4-7, TTX significantly altered spiking pattern, converting all but one neuron to transient spiking (*χ*^2^=20.05, p<0.0001), and it significantly reduced firing rate (F_1,132_=43.157, p<0.001, n=23). Only neurons with repetitive spiking at baseline are included in the firing rate plot. **(B)** At DIV0, TTX did not affect rheobase (Z_24_=1.129, p=0.265, Wilcoxon rank test) but did reduce spike height (T_24_=3.092, p=0.005, paired t-test). At DIV4-7, TTX increased rheobase (Z_28_=4.681, p<0.001, Wilcoxon rank test) and dramatically reduced spike height (T_28_=20.333, p<0.001, paired t-test). Notably, neurons at DIV0 and DIV4-7 did not differ in their baseline rheobase (U=316, p=0.425, Mann-Whitney test) or spike height (T_52_=0.322, p=0.749, t-test). **(C)** Neurons at DIV0 and DIV4-7 differed in their total capacitance (T_52_=6.728, p<0.001, t-test) and leak conductance density (U=216, p=0.011, Mann-Whitney test) but not in their resting membrane potential (T_52_=1.668, p=0.101, t-test). **(D)** Sample voltage clamp recordings with command voltage stepped from -85 mV to +15 mV in 5 mV increments, before and after TTX. Sodium current was not significantly reduced by TTX at DIV0 (F_1,72_=3.585, p=0.107, two-way RM ANOVA; n=7 neurons) but was completely abolished by TTX at DIV4-7 (F_1,108_=33.526, p<0.001; n=10 neurons). *, p<0.05; **, p<0.01; Student-Newman-Keuls post-hoc tests in A and D. Traces labeled “TTX-sensitive current” represent the difference between current measured at baseline and after TTX, as determined by subtracting responses to the same voltage step under different pharmacological conditions.

### Different Na_V_ channel subtypes control nociceptor excitability at DIV0 and DIV4-7

Next, we sought to identify the Na_V_ subtype responsible for repetitive spiking at each time point, starting with DIV0. Of the TTX-R Na_V_ channels expressed by nociceptors, Na_V_1.8 has been implicated in repetitive spiking (47, 48). We measured sodium current in voltage clamp before and after applying the Na_V_1.8-selective inhibitor PF-01247324 (PF-24) (50). At DIV0, 1 µM PF-24 abolished most of the sodium current (**Fig. 2A**). The PF-24-sensitive current had slow inactivation kinetics, like the TTX-R current and unlike the fast TTX-S current in Figure 1D, and consistent with previous descriptions of Na_V_1.8 (51). A different Na_v_1.8 antagonist, A-803467, had similar effects (**Fig. 2 – figure supplement 1**). In current clamp, PF-24 converted 7 of 8 repetitively spiking neurons to transient spiking and significantly reduced evoked spiking (**Fig. 2B**). It also increased rheobase and decreased spike height but did not affect resting membrane potential (**Fig. 2C**). PF-24 had negligible effects when tested at DIV4-7 (**Fig. 2 – figure supplement 2**). These results show that Na_V_1.8 is the predominant Na_V_ subtype at DIV0 and is necessary for repetitive spiking at that time point. To test the sufficiency of Na_V_1.8 to produce repetitive spiking, we tuned a single-compartment, conductance-based model neuron (see Methods) to reproduce DIV0 data described above. In this DIV0 model, inclusion of Na_V_1.8 conductance was sufficient to generate repetitive spiking (**Fig. 2D** left). The necessity of Na_V_1.8 for repetitive spiking at DIV0 was also recapitulated: 85% reduction in the Na_V_1.8 conductance converted spiking from repetitive to transient (**Fig. 2D** and **Supplementary Table 1**).

**Figure 2.**
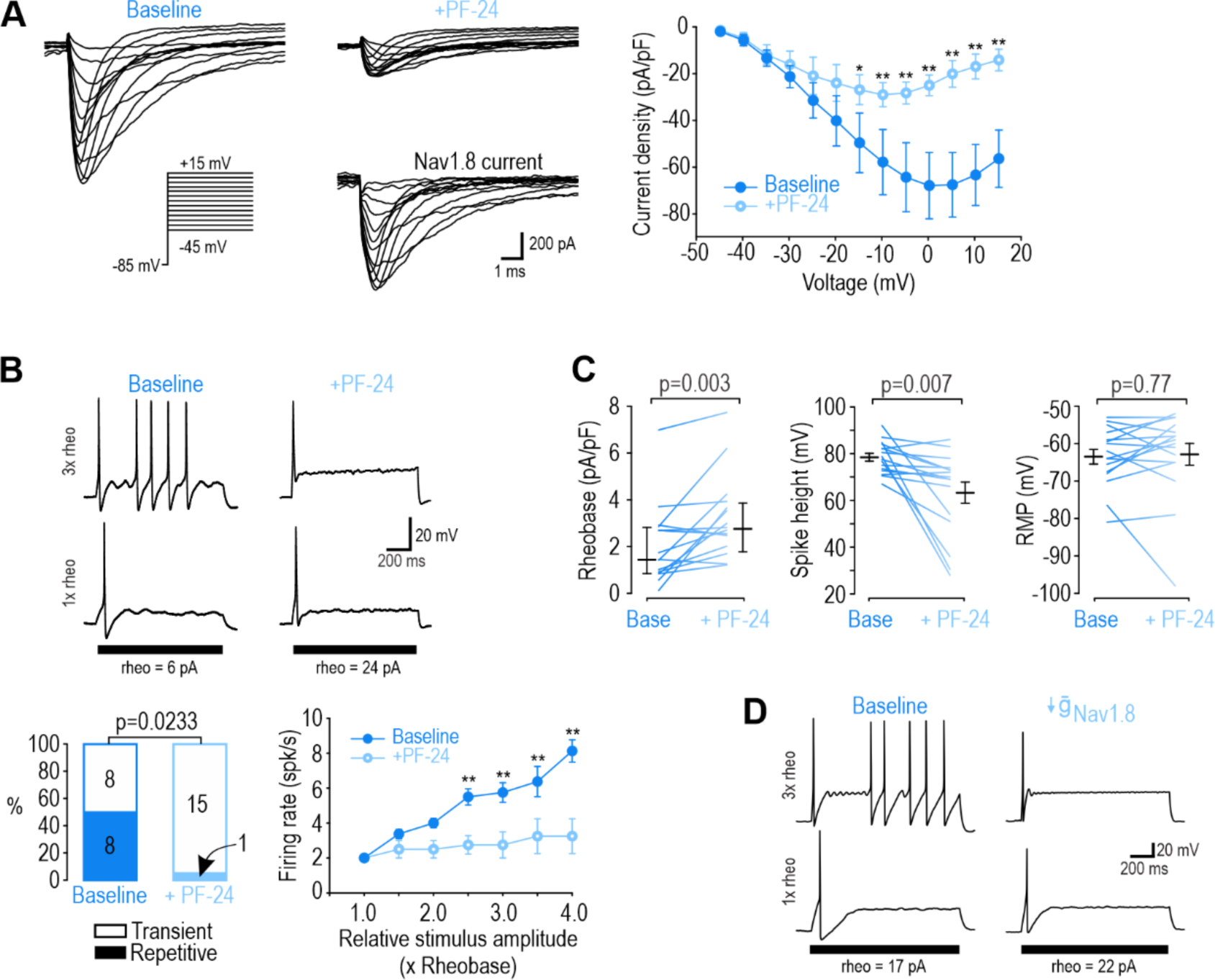
Na_V_1.8 is necessary for repetitive spiking at DIV0. **(A)** Sample voltage clamp recordings show that sodium current was almost completely abolished by the Na_V_1.8 inhibitor PF-24 (1 µM). Peak current was significantly reduced by PF-24 (F_1,72_=12.651, p<0.012, two-way RM ANOVA; n=7). Traces labeled “Na_V_1.8 current” represent the difference between current measured at baseline and after PF-24, as determined by subtraction. Another Na_V_1.8 inhibitor, A-803467, had a similar effect (see Fig. 2 – figure supplement 1). **(B)** PF-24 significantly altered spiking pattern (χ^2^=5.14, p=0.0233, McNemar test) and reduced firing rate (F_1,42_=11.946, p=0.011, two-way RM ANOVA; n=8). **(C)** PF-24 significantly increased rheobase (Z_15_=2.783, p=0.003, Wilcoxon rank test) and reduced spike height (T_15_=3.151, p=0.007, paired t-test) but did not affect resting membrane potential (T_15_=0.304, p=0.765, paired t-test). PF-24 had limited effects at DIV4-7 (Fig 2 – figure supplement 2). **(D)** A computational model reproduced the effect of Na_V_1.8 on spiking pattern (also see Supplementary Table 1). The PF-24 effect was simulated as a ∼85% reduction in Na_V_1.8 (*ḡ*_Nav1.8_= 4mS/cm2). *, p<0.05; **; p<0.01; Student-Newman-Keuls post-hoc tests in A and B.

Next, we sought to identify the Na_V_ subtype responsible for repetitive spiking at DIV4-7 using PF-05089771 (PF-71) to inhibit Na_V_1.7 (40, 52) and ICA-121431 (ICA) to inhibit Na_V_1.1/1.3 (53, 54). Since Na_V_1.1 is expressed mostly in medium-diameter (Aδ) neurons (55) whereas Na_V_1.3 is known to be upregulated in C fibers after injury (for review, see 56), we ascribe the ICA effect to blockade of Na_V_1.3. In voltage clamp, sodium current was significantly reduced by 30 nM PF-71, and most of the remaining current was blocked by 1 µM ICA (**Fig. 3A**). In current clamp, each inhibitor (applied separately) converted a significant proportion of neurons to transient spiking and significantly reduced firing rate (**Fig. 3B**). This argues that Na_V_1.7 and Na_V_1.3 are both necessary for repetitive spiking at DIV4-7. Inhibiting Na_V_1.7 increased rheobase, unlike inhibiting Na_V_1.3, and caused a stronger reduction in spike height (**Fig. 3C**). Neither affected resting membrane potential. These results show that Na_V_1.7 is the predominant Na_V_ subtype at DIV4-7, but not the only one. PF-71 had negligible effects when tested at DIV0 (**Fig. 3 – figure supplement 1)**. We re-tuned our computational model to reproduce DIV4-7 data, with both Na_V_1.7 and Na_V_1.3 being required to produce repetitive spiking, meaning neither channel is individually sufficient (**Fig. 3D** and **Supplementary Table 1**). That said, inserting a higher density of either Na_V_1.7 or Na_V_1.3 could produce repetitive spiking in the absence of the other subtype (**Fig. 3 – figure supplement 2)**, consistent with Na_V_1.7 and Na_V_1.3 also being interchangeable.

**Figure 3.**
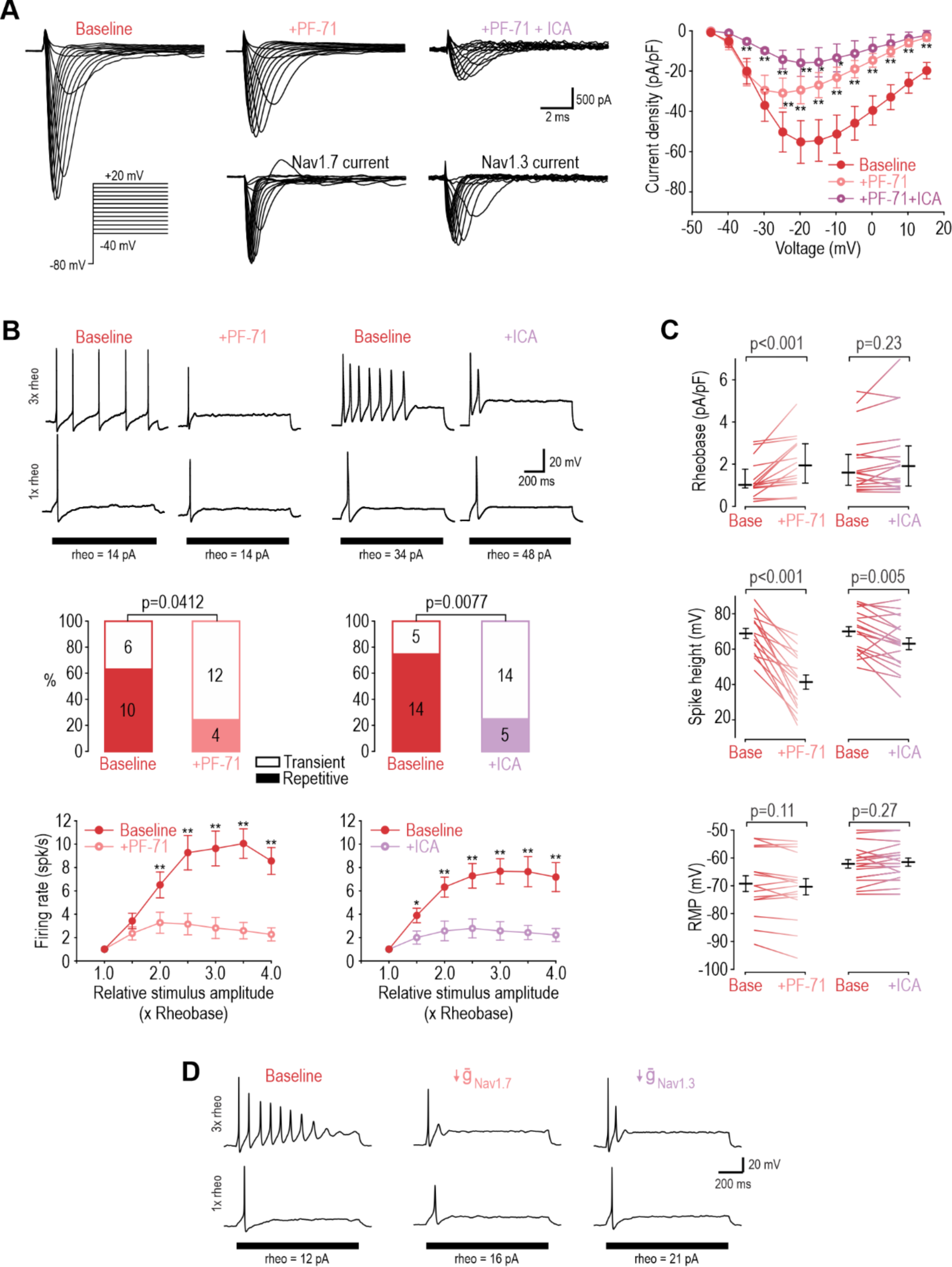
Na_V_1.3 and Na_V_1.7 are necessary for repetitive spiking at DIV4-7. **(A)** Sample voltage clamp recordings show that sodium current was reduced by the Na_V_1.7 inhibitor PF-71 (30 nM) and by the Na**_V_**1.1/1.3 inhibitor ICA (1 µM). Peak current was significantly reduced by PF-71 and ICA (F_2,192_=26.361, p<0.001, two-way RM ANOVA; n=9). Traces labeled “Na_V_1.7 current” and “Na_V_1.3 current” represent the difference between current measured at baseline and after PF-72 and ICA, respectively, as determined by subtraction. **(B)** PF-71 and ICA both significantly altered spiking pattern (χ^2^=4.17, p=0.041 and χ^2^ =7.11, p=0.0077, respectively, McNemar tests) and significantly reduced firing rate (F_1,54_=40.659, p<0.001, n=10 and F_1,78_=35.156, p<0.001, n=14, respectively, two-way RM ANOVAs). **(C)** PF-71 significantly increased rheobase (Z_18_=3.464, p<0.001, Wilcoxon rank test) and decreased spike height (T_18_=7.946, p<0.001, paired t-test). ICA did not significantly alter rheobase (Z_18_=1.248, p=0.225) but did reduce spike height (T_18_=3.243, p=0.005). Neither drug affected resting membrane potential (T_15_=1.681, p=0.113 for PF-71; T_18_=-1.132, p=0.272 for ICA, paired t-test). PF-71 had negligible effects at DIV0 (Fig. 3 – figure supplement 1). **(D)** A computational model reproduced the combined effects of Na_V_1.3 and Na_V_1.7 on spiking pattern (also see Supplementary Table 1 and Fig. 3 – figure supplement 2). PF-71 effect was simulated as a 70% reduction in Na_V_1.7 (*ḡ*_Nav1.7_= 10.5 mS/cm^2^). ICA effect was simulated as a 90% reduction in Na_V_1.3 (*ḡ*_Nav1.3_’= 0.035 mS/cm^2^). *, p<0.05; **, p<0.01; Student-Newman-Keuls post-hoc tests in A and B.

### Acutely interchanging Na_V_ subtypes does not affect spiking pattern

The ability of Na_V_1.3, Na_V_1.7 and Na_V_1.8 to each encourage repetitive spiking is seemingly inconsistent with the common view that each Na_V_ subtype contributes selectively to a different phase of the spike (for example, Figure 3 in ref. 56). If Na_V_1.8 were to activate exclusively at suprathreshold voltages, it could not initiate spikes and a different perithreshold-activating Na_V_ channel would be needed, which is clearly inconsistent with our data. To verify that Na_V_1.7 and Na_V_1.8 currents are each sufficient to produce repetitive spiking, we tested whether the Na_V_1.8 current necessary for spiking in our DIV0 computer model could be replaced with Na_V_1.7, and whether the Na_V_1.7 current necessary for spiking in our DIV4-7 computer model could be replaced with Na_V_1.8. In both cases, repetitive spiking was restored after inserting the alternate current (**Fig. 4A**). We then proceeded with equivalent experiments in real neurons, inhibiting Na_V_1.8 with PF-24 on DIV0 or Na_V_1.7 with PF-71 on DIV4-7, and then introducing the alternate channel virtually using dynamic clamp (see *Methods*). The replacement was successful in all neurons tested (**Fig. 4B**). Inserting virtual Na_V_1.8 after inhibiting native Na_V_1.8 also restored repetitive spiking, and likewise for Na_V_1.7 (**Fig. 4 – figure supplement 1**), verifying that our virtual channels were equivalent to the native channels we aimed to replace. Apart from maximal conductance density, which was titrated in each neuron, all other parameters used for dynamic clamp were identical to simulations. The success of dynamic clamp experiments helps validate our computational models insofar as virtual Na_V_1.7 and Na_V_1.8 currents interacted appropriately with native currents to produce repetitive spiking in real neurons, the same way they interact with other simulated currents in the model neuron. Please note that tests reported In **Figure 4B** involve replacing a native channel with a different virtual channel (e.g. native Na_V_1.8 replaced with virtual Na_V_1.7) whereas tests reported in **Figure 4 – figure supplement 1** involve replacing a native channel with the equivalent virtual channel (e.g. native Na_V_1.8 replaced with virtual Na_V_1.8); the former demonstrates that Na_V_ subtypes are interchangeable whereas the latter serves as a positive control.

**Figure 4.**
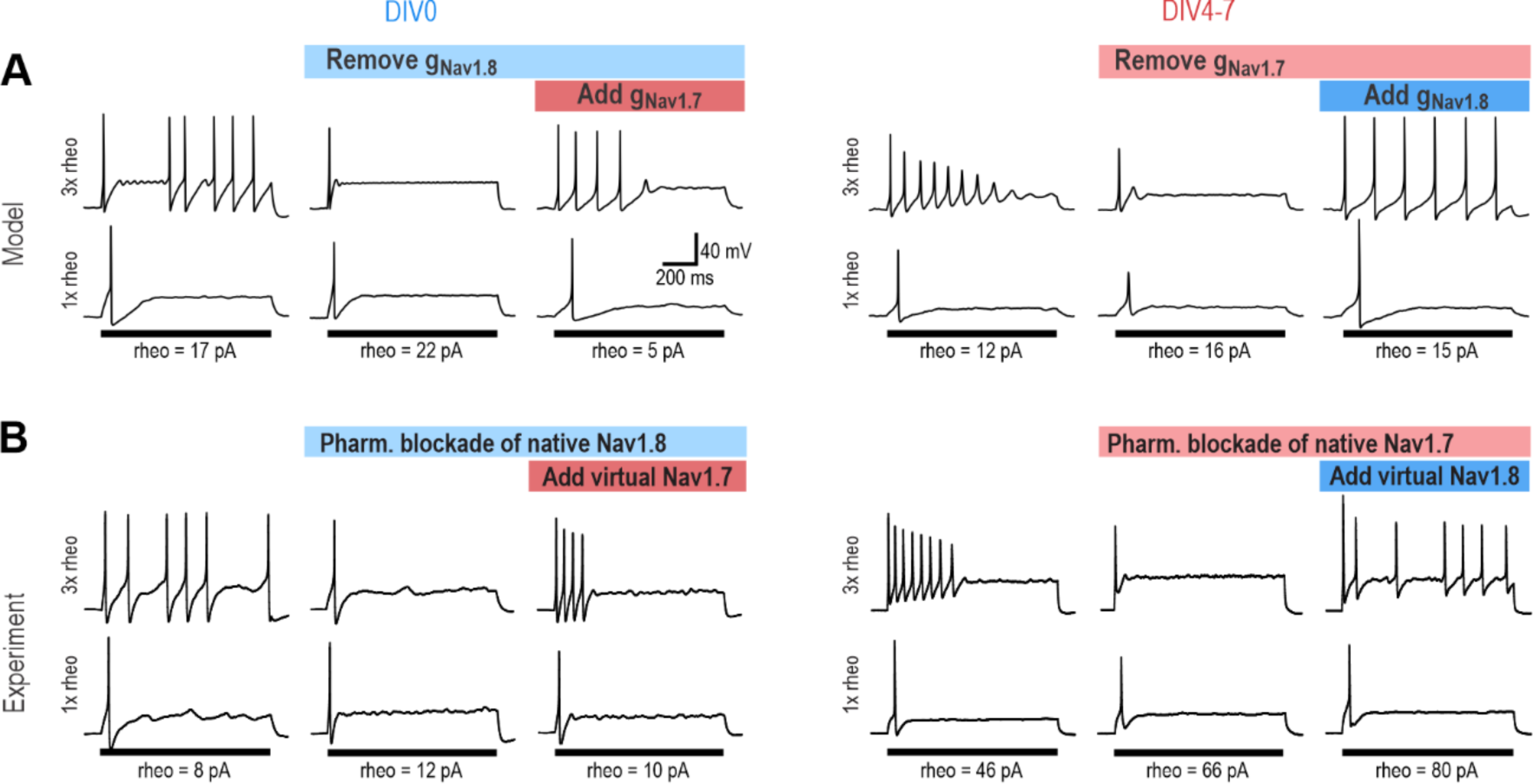
Na_V_1.7 and Na_V_1.8 are each sufficient to produce repetitive spiking in DIV0 and DIV4-7 neurons. **(A)** The computational model predicts that the Na_V_1.8 conductance, which is “necessary” for repetitive spiking at DIV0 can, in principle, be replaced by Na_V_1.7 (left), and vice versa at DIV4-7 (right). **(B)** Replacement experiments involved inhibiting native channels pharmacologically and then introducing virtual conductances using dynamic clamp. At DIV0 (left), inhibiting native Na_V_1.8 (with PF-24) converted neurons to transient spiking, but introducing virtual Na_V_1.7 reverted neurons to repetitive spiking (in 3 of 3 neurons tested). At DIV4-7, inhibiting native Na_V_1.7 (with PF-71) converted the neuron to transient spiking, but introducing virtual Na_V_1.8 reverted neurons to repetitive spiking (in 4 of 4 neurons tested). Repetitive spiking was likewise restored by replacing the blocked native channel with the corresponding virtual channel (Fig. 4 **– figure supplement 1**). Parameters for virtual channels were identical to simulations except for the maximal conductance density, which was titrated in each cell.

With the model neurons thus validated, we used simulations to infer Na_V_1.7 and Na_V_1.8 currents during different phases of the spike (**Fig. 5A-D**). Since inward (depolarizing) current at voltages just below spike threshold is critical for spike initiation (57), we sought to identify which Na_V_ contributes to the subthreshold current. In the DIV0 model (**Fig. 5A,B**), subthreshold inward current was mediated mostly by Na_V_1.7 during the first spike (left) but by Na_V_1.8 during the second and all subsequent spikes (right). We interpret this to mean that the first spike is initiated using Na_V_1.7 whereas all subsequent spikes are initiated using Na_V_1.8. This is explained by the small Na_V_1.7 conductance at DIV0 quickly inactivating during the first spike and remaining inactive during subsequent spikes (**Fig. 5 – figure supplement 1A**). This is consistent with experimental results, where repetitive spiking at DIV0 was unaffected by inhibiting Na_V_1.7 (see Fig. 1 and Fig. 3 – figure supplement 1) but was prevented by inhibiting Na_V_1.8 (see Fig. 2). Inactivation of Na_V_1.7 after the first spike was reflected by an increase in voltage threshold between the first and second spike in the model (**Fig. 5A**), which prompted us to check if the same increase was evident experimentally, which it was (**Fig. 5E**). This unanticipated simulation result also predicted that TTX should affect the voltage threshold of the first spike in DIV0 neurons despite not having other notable effects (see Fig. 1); as predicted, TTX caused a significant depolarizing shift in voltage threshold at DIV0 (**Fig. 5 – figure supplement 2)**, further validating our model. In the DIV4-7 model **(Fig. 5C,D)**, subthreshold inward current was mediated by Na_V_1.7 and Na_V_1.3 during the first spike (left) and during all subsequent spikes (right), with Na_V_1.8 contributing little. Even though inactivation reduced Na_V_1.7 and Na_V_1.3 current after the first spike (**Fig. 5 – figure supplement 1B)**, those channels nonetheless provided sufficient inward current to support repetitive spiking at DIV4-7. Inactivation at DIV4-7 was reflected, however, in a combination of higher threshold and lower spike overshoot for the second spike, both in the model (**Fig. 5C,D**) and in experiments (**Fig. 5E**).

**Figure 5.**
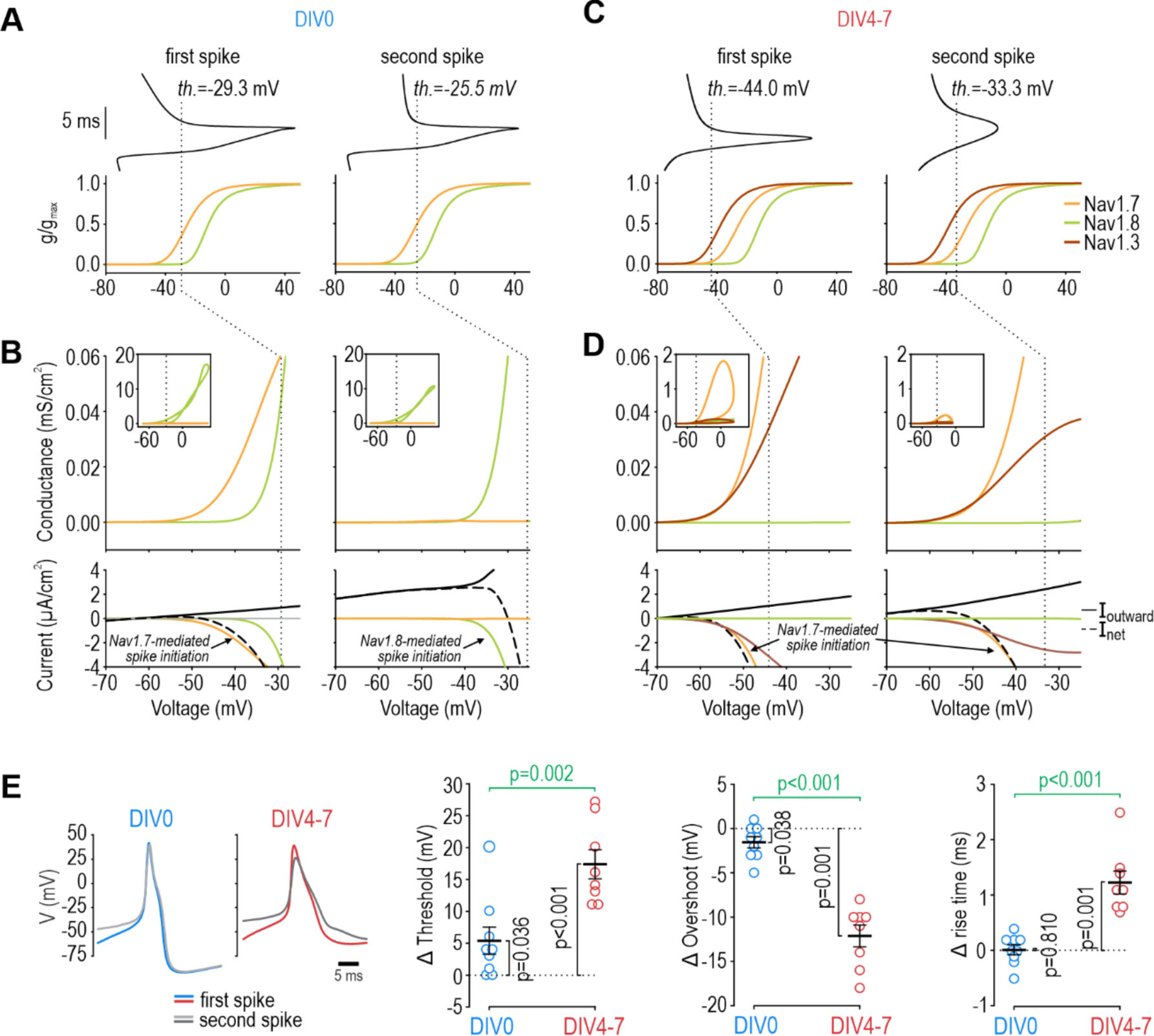
Contribution of Na_V_1.7 and Na_V_1.8 to spike initiation in DIV0 and DIV4-7 neurons. **(A)** Voltage (top) for first (left) and second (right) spikes in the DIV0 model aligned with voltage activation curves for each Na_V_ subtype (bottom). Dashed line shows voltage threshold (defined as V where dV/dt reaches 5 mV/ms). **(B)** Conductance plotted against voltage to create a phase portrait (top) showing Na_V_ conductance at different phases of the spike. Inset shows full voltage range; main graph zooms in on voltages near threshold. Bottom plots show current plotted over the same voltage range. Whereas Na_V_1.7 (orange) mediated nearly all perithreshold inward current for the first spike, voltage threshold increased – because Na_V_1.7 inactivated (Fig. 5 – figure supplement 1) – and Na_V_1.8 (green) mediated nearly all perithreshold inward current for the second spike. The unexpected contribution of Na_V_1.7 to the first spike correctly predicted that TTX increases voltage threshold in DIV0 neurons (Fig. 5 – figure supplement 2). (**C, D**) In the DIV4-7 model, Na_V_1.7 (orange) and Na_V_1.3 (maroon) contributed to initiation of all spikes whereas the contribution of Na_V_1.8 was negligible (due entirely to its low expression level). **(E)** Sample experimental traces showing differences in the first (blue/red) and second (grey) spikes at DIV0 and DIV4-7. Plots summarize differences (Δ) in threshold, overshoot potential, and spike rise time between 1^st^ and 2^nd^ spikes during repetitive spiking evoked by current injection. At DIV0, the 1^st^ and 2^nd^ spikes differ significantly in their threshold (T_8_=2.522, p=0.036, one-sample t-test) and overshoot (T_8_=0.038, p=0.038) but not rise time (T_8_=0.249, p=0.810). At DIV4-7, the 1^st^ and 2^nd^ spikes differ in all measures (threshold: T_7_=7.613, p<0.001; overshoot: T_7_=-9.849, p<0.001; rise time: T_7_=5.979, p<0.001). Statistical results (green) show that differences between 1^st^ and 2^nd^ spike at DIV4-7 are significantly larger than differences at DIV0 (threshold: T_15_=-3.847, p=0.002; overshoot: T_15_=7.922, p<0.001; rise time: T_15_=-5.617, p<0.001, unpaired t-tests), consistent with our computational model.

These results demonstrate that each Na_V_ subtype does not contribute exclusively to a particular phase of the spike, and nor is each spike phase mediated exclusively by a particular Na_V_ subtype. Instead, each Na_V_ subtype contributes preferentially to a different spike phase depending on its voltage-dependency and on the conductance densities and inactivation status of other Na_V_ subtypes; for instance, Na_V_1.8 is often said to activate only at suprathreshold voltages, during the upswing of the spike, after the spike is initiated by Na_V_1.7; but if Na_V_1.7 is absent or inactivated, voltage threshold shifts into the Na_V_1.8 activation range, thus enabling Na_V_1.8 to activate at voltages that are now subthreshold. Indeed, a subtype’s contribution can shift rapidly (because of channel inactivation) or slowly (because of changes in conductance density; see below).

### Changes in Na_V_ subtype expression between DIV0 and DIV4-7

Next, we sought to identify the basis for the slow shift in which Na_V_ subtype controls nociceptor excitability. **Figure 6A** shows mRNA levels for Na_V_1.7 and Na_V_1.8 relative to a housekeeping gene (left) and to each other (right). Na_V_1.7 mRNA levels exceeded Na_V_1.8 mRNA levels at both DIV0 and DIV7. Both decreased between DIV0 and DIV7, but Na_V_1.8 more so, resulting in a significant decrease in the Na_V_1.8:Na_V_1.7 mRNA ratio. This pattern is consistent with the reduced role of Na_V_1.8 at DIV4-7 but is inconsistent with the negligible role of Na_V_1.7 at DIV0; specifically, we expected Na_V_1.7 mRNA levels to increase between DIV0 and DIV7. Next, we investigated if functional changes were better reflected by changes in protein levels. Immunofluorescence for Na_V_1.8 was higher than for Na_V_1.7 at DIV0, and that ratio reversed at DIV7 (**Fig. 6B**), consistent with functional changes. Moreover, cercosporamide (10 µM), a potent inhibitor of the eukaryotic translation Initiation Factor 4E (eIF4E), significantly mitigated the decrease in Na_V_1.8 immunofluorescence and the increase in Na_V_1.7 immunofluorescence when applied to cultured neurons for 24 or 120 hours prior to measurements on DIV5 (**Fig. 6C**). Beyond showing that their mRNA levels do not correlate well with Na_V_ contributions to nociceptor excitability, reminiscent of some previous work (e.g. (58)), these results suggest that translational regulation is crucial, though membrane trafficking and other downstream processes likely also contribute (59, 60). Specifically, our results suggest that there is not enough pre-existing Na_V_1.7 channels that trafficking those channels to the membrane can explain observed functional changes; instead, synthesis of new Na_V_1.7 protein from existing Na_V_1.7 mRNA is involved, but that does not rule out changes in how Na_V_1.7 protein is handled or why Nav1.8 decreases. Further investigation is required to explore those mechanisms.

**Figure 6.**
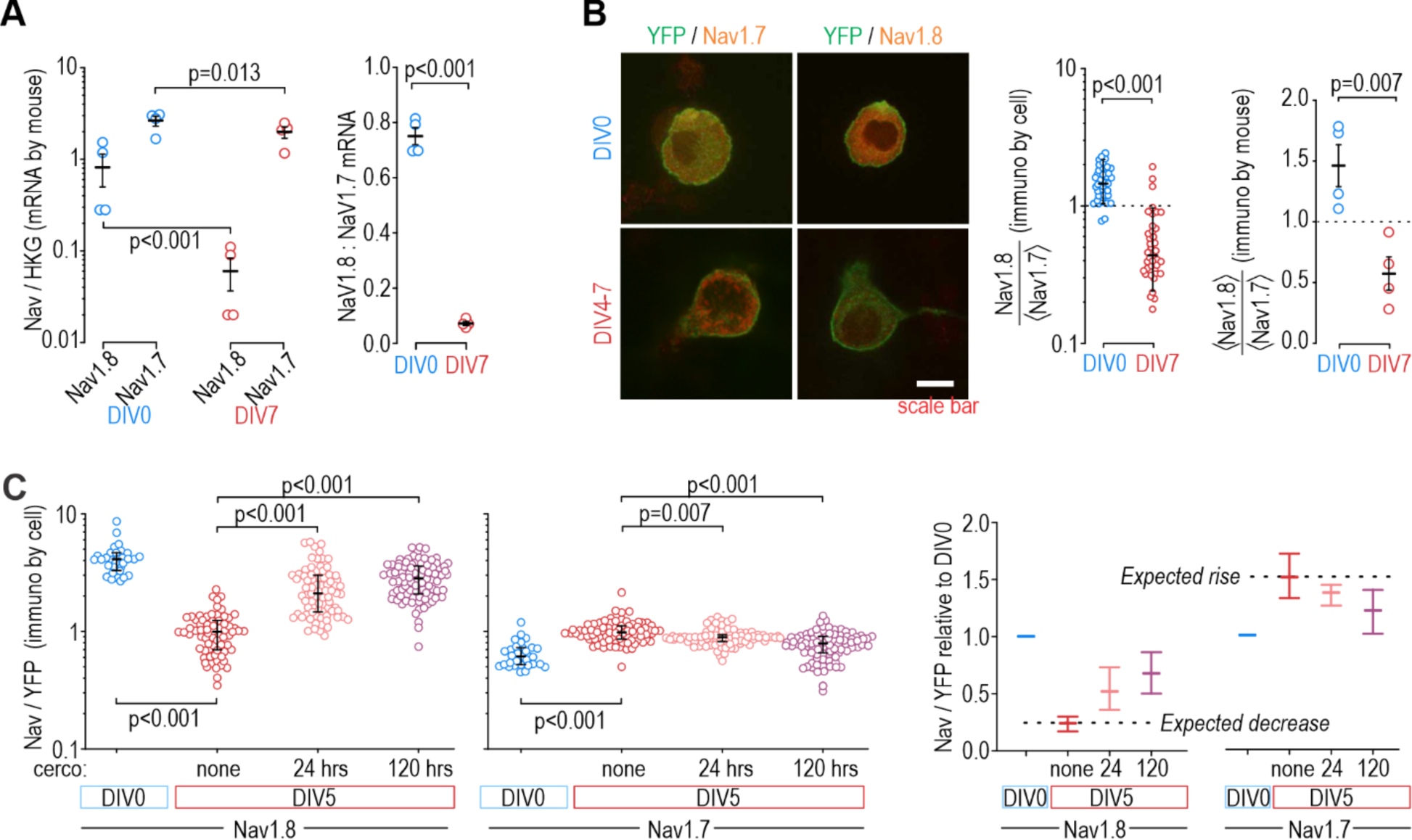
Protein levels, but not mRNA, reflect functional contributions of Na_V_ subtypes at DIV0 and DIV7. **(A)** Both Na_V_1.8 and Na_V_1.7 mRNA levels (relative to a housekeeping gene (HKG), see Methods) decreased significantly between DIV0 and DIV4-7 (factor 1: time, F_1,12_=56.677, p<0.001, factor 2: subtype, F_1,12_=17.952, p=0.001, two-way ANOVA and Student-Newman-Keuls post-hoc tests on log transformed data, n=4 mice per time point) but more so for Na_V_1.8 than for Na_V_1.7 (interaction: time x subtype, F_1,12_= 11.455, p=0.005). The differential reduction yielded a significantly higher Na_V_1.8: Na_V_1.7 ratio at DIV0 than at DIV7 (T_6_=21.375, p<0.001, unpaired t-test) but the increasing functional contribution of Na_V_1.7 between DIV0 and DIV4-7 remains unaccounted for. **(B)** Immunoreactivity (IR) for Na_V_1.8 protein exceeded Na_V_1.7-IR at DIV0, but the opposite was true on DIV4-7, consistent with the functional contribution of each subtype. Na_V_-IR was measured relative to YFP intensity in the same cell, and then each cell’s Na_V_1.8:YFP ratio was considered relative to the average Na_V_1.7:YFP ratio in the co-processed coverslip (left) or average Na_V_1.8:YFP ratio was considered relative to the average Na_V_1.7:YFP ratio in the same animal (right). Ratios were >1 at DIV0 but decreased significantly at DIV4-7 (U=78, p<0.001, n=37 for DIV0, n=40 for DIV4-7, Mann-Whitney test (left) and T_6_=4.046, p=0.007, unpaired t-test (right)). **(C)** Chronically applied cercosporamide (10 µM) mitigated changes in Na_V_1.8- and Na_V_1.7-IR at DIV5 (Na_V_1.8: H_3_=157.95, p<0.001; Na_V_1.7: H_3_=80.662, p<0.001; One-way ANOVA on ranks, Dunn’s post-hoc tests, p<0.05 for all pairs). Data are summarized as median±quartile. Panel on the right shows data normalized to baseline (DIV0) to emphasize relative changes. N = 3 experiments.

### Analgesic efficacy of subtype-selective drugs depends on which Na_V_ controls nociceptor excitability

If a Na_V_1.7-selective inhibitor mediates analgesia by modulating nociceptor excitability, its analgesic efficacy hinges on nociceptor excitability being controlled by Na_V_1.7. Accordingly, we predicted that the Na_V_1.7-selective inhibitor PF-71 would have little if any effect on paw withdrawal under normal conditions, when Na_V_1.8 controls nociceptor excitability (Fig. 2 and Fig. 3 – figure supplement 1), but would be effective if Na_V_1.7 took over control. Inflammation increases Na_V_1.7 channel trafficking and membrane expression (61-64). To test if inflammation increased Na_V_1.7’s influence on nociceptor excitability, we recorded neurons acutely dissociated (DIV0) from DRGs of mice whose hind paw was injected with CFA three days prior. Sample traces in Figure 7A show that inflammation caused nociceptors to become much more variable in their reliance of specific Na_V_ subtypes, presumably because neurons innervating the inflamed paw experience greater effects of inflammation than neurons innervating other, non-inflamed sites. Specifically, application of PF-71 to inhibit Na_V_1.7 converted 5 of 12 (42%) CFA neurons to transient spiking vs 0 of 9 (0%) of control neurons, which is a significantly higher proportion (p = 0.0451, Fisher exact test, **Fig. 7B, left**), whereas subsequent application of PF-24 to inhibit Na_V_1.8 converted only 1 of 7 (14%) of the remaining repetitive spiking CFA neurons to transient spiking vs 7 of 8 (88%) of control neurons, which is a significantly lower proportion (p = 0.001). PF-71 also significantly affected resting membrane potential, rheobase, and spike height after CFA (**Fig. 7C**), unlike in control neurons (see Fig. 3 – figure supplement 1).

**Figure 7.**
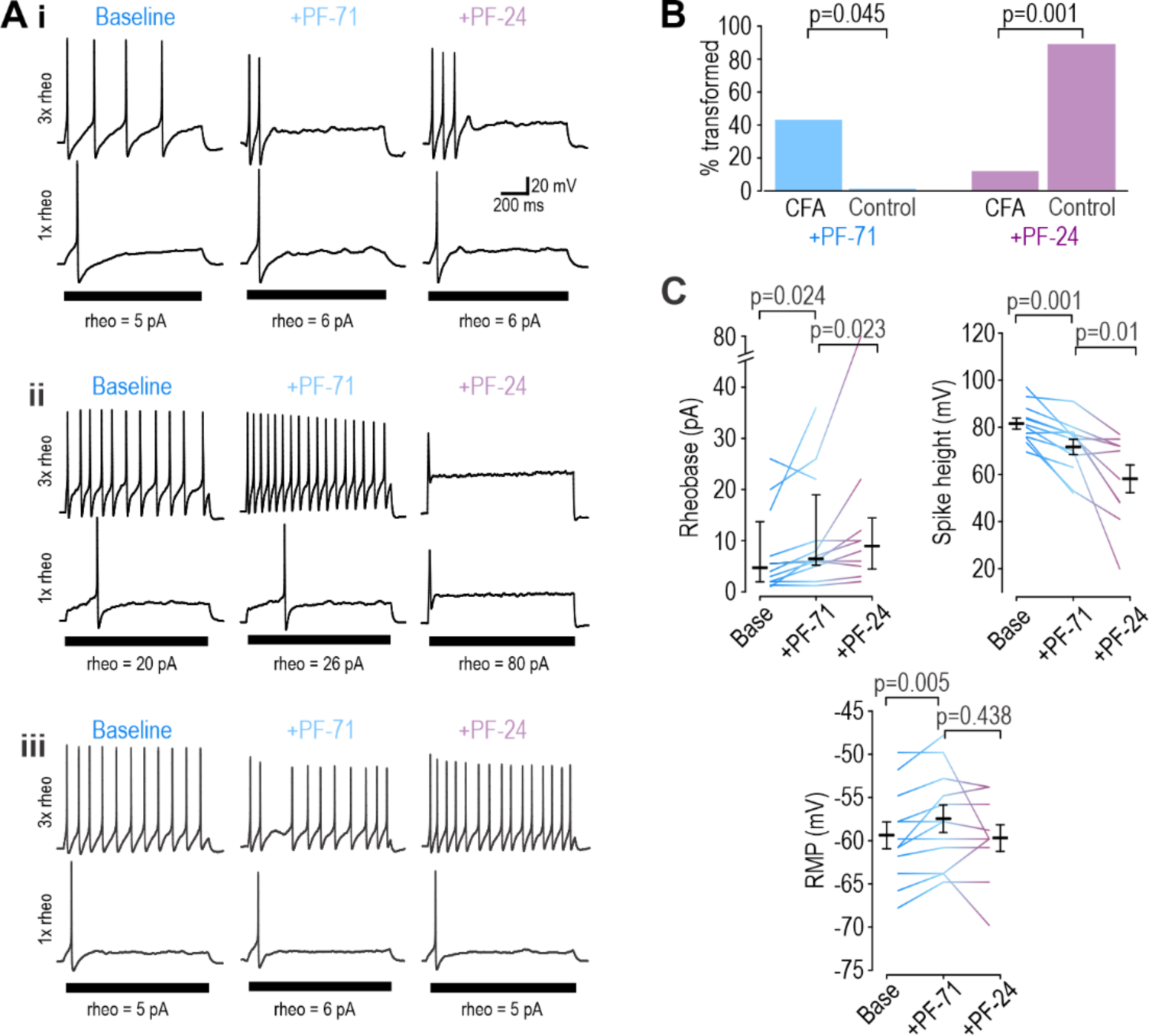
Inflammation alters Na_V_ subtype contribution to nociceptor excitability. (**A**) Sample responses in DIV0 neurons from mice injected with CFA three days earlier. In 12 cells tested, PF-71 converted 5 neurons to transient spiking (**i**), encouraged repetitive spiking in 4 neurons (**ii**), and had no effect in 3 neurons (**iii**), thus highlighting increased heterogeneity after CFA. (**B**) At DIV0, the effect of PF-71 differed significantly between CFA and control neurons, converting 42% (5 of 12) CFA neurons from repetitive to transient spiking vs 0% (0 of 9) control neurons (p=0.0451, Fisher Exact test). Applying PF-24 to neurons that continued to spike repetitively after PF-71 had little effect on CFA neurons, converting only 13% (1 of 7) of CFA neurons vs 88% (7 of 8) of control neurons (p=0.001, Fisher Exact test). Together these results argue that Na_V_1.7 contributes more and Na_V_1.8 contributes less to nociceptor excitability after inflammation. **(C)** At DIV0, PF-71 significantly increased resting membrane potential (T_11_=-3.530, p=0.005, paired t-test) and rheobase (Z_11_=2.186, p=0.024, Wilcoxon rank test), and significantly decreased spike height (T_11_=4.413, p=0.001, paired t-test) in CFA neurons. Further addition of PF-24 significantly changed rheobase (Z_9_=2.176, p=0.023, Wilcoxon rank test) and spike height (T_9_=3.237, p=0.01, paired t-test) but did not affect resting membrane potential (T_9_=1.049, p=0.321, paired t-test).

Results above confirm that Na_V_1.7 takes on greater responsibility for nociceptor excitability after inflammation, which in turn predicts that PF-71 should reduce pain after inflammation but not under control conditions. As predicted, PF-71 significantly reduced thermal (**Fig. 8A**) and tactile (**Fig. 8B**) sensitivity in CFA-inflamed mice without having any effect in control mice. These results show that the inflammation-induced shift in Na_V_ subtype expression, despite being variable (see Fig. 7), is sufficient to cause a measurable change in drug efficacy assessed in vivo. Consistent with this, epigenetic repression of Na_V_1.7 prevents/reverses hypersensitivity in inflamed and neuropathic mice without causing hyposensitivity in naïve mice (65). This is unlike genetic deletion of Na_V_1.7, which reduces thermal and tactile sensitivity in naïve mice (27), and with loss-of-function mutations in Na_V_1.7 that abolish pain in humans (17). These inconsistencies rekindle concerns whether Na_V_1.7 mutations, unlike pharmacological interventions, affect pain through mechanisms other than modulation of nociceptor excitability. Pharmacological reversal of hypersensitivity in chronic pain conditions (when Na_V_1.7 is pathologically upregulated) without reducing normal nociceptive pain is clinically desirable, but this hinges on nociceptor hyperexcitability being Na_V_1.7-dependent, which may be true of some but not all chronic pain conditions, or in only a subset of patients (66) (see Discussion).

**Figure 8.**
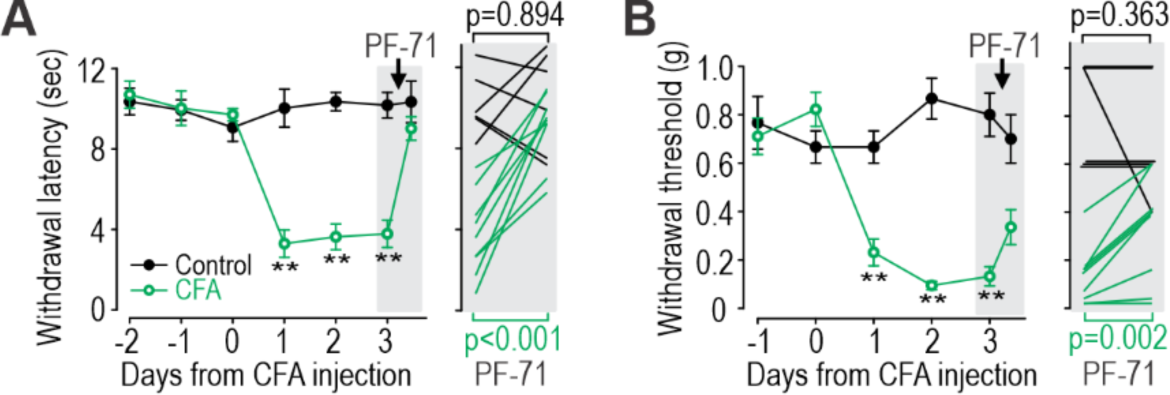
Inflammation-induced change in Na_V_ subtype contribution impacts analgesic efficacy of PF-71. **(A)** CFA significantly increased thermal sensitivity (F_5,65_=19.556, p<0.001, two-way RM ANOVA). PF-71 significantly decreased thermal sensitivity in mice injected three days prior with CFA (T_8_=-7.296, p<0.001; paired t-test) but had no effect in naïve mice (T_5_=-0.141, p=0.894). **(B)** CFA significantly increased mechanical sensitivity (F_4,52_=16.786, p<0.001). PF-71 significantly decreased tactile sensitivity in mice injected three days prior with CFA (T_8_=-4.341, p=0.002) but had no effect in naive mice (T_5_=1.000, p=0.363). Insets in both panels show values for each animal before and 2 hours after PF-71 injection. *, p<0.05; **, p<0.01; Student-Newman-Keuls post-hoc tests.

## DISCUSSION

Our results show that nociceptors can achieve similar excitability using different Na_V_ channels. Whereas repetitive spiking depends on Na_V_1.8 shortly after dissociation (Fig. 2) and presumably under normal conditions in vivo, responsibility shifts to Na_V_1.7 and Na_V_1.3 after a few days in vitro (Fig. 3). This is due to translationally regulated changes in Na_V_ expression (Fig. 6). Inflammation causes a similar shift in vivo (Fig. 7). Importantly, acutely inhibiting a particular Na_V_ is consequential (analgesic) only if that subtype is responsible for nociceptor excitability (Fig. 8). This may explain why Na_V_1.7-selective drugs have not performed well in clinical trials (see Introduction) – because Na_V_1.7 is not always necessary for nociceptor excitability depending on the expression level of Na_V_1.7 and other Na_V_ subtypes. Faster processes like channel inactivation also affect their relative contribution (Fig. 5). These observations demonstrate the variable contribution of different Na_V_ subtypes to nociceptor excitability. When unaccounted for, such variability can lead to inconsistencies at the root of poor reproducibility and translatability.

Although we have focussed here on Na_V_ channels, numerous other channels are likely to change during culturing and in response to subtler, more clinically relevant in vivo insults like inflammation. Those changes occur through diverse mechanisms. Our results argue that transcriptional changes in Na_V_1.7 do not account for its upregulation between DIV0 and DIV4-7, but transcriptional changes in other genes might nonetheless be important. And though our data implicate translational regulation, much more work is needed to work out the details. That work should consider the other ion channels and associated proteins (e.g. beta subunits) that interact with Na_V_ channels, either physically or via mutual effects on membrane potential. In a degenerate system, one should ideally consider all the components contributing to the process of interest, but perfect is the enemy of good, which is to say that degeneracy should still be considered even with incomplete understanding of the system. Indeed, the interchangeability of Na_V_ subtypes demonstrated here may help explain why subtype-specific drugs are not reliably effective again pain, even if myriad other still unidentified changes are also taking place.

Contrary to the view that certain ion channels are uniquely responsible for certain aspects of neuronal function, neurons use diverse ion channel combinations to achieve similar function (67, 68). This degeneracy is crucial for enabling excitability and other aspects of neuron physiology to be homeostatically regulated by adjusting ion channels in response to perturbations (69-71). Degeneracy also enables pathological changes in different ion channels to produce equivalent hyperexcitability (72). This is important insofar as similar excitability may belie differences in the underlying ion channels – differences that may render a neuron susceptible or impervious to a drug depending on the functional necessity of the targeted ion channel in that neuron. This is precisely what our data demonstrate in nociceptors. Similar observations have been made in substantia nigra neurons, whose pacemaker activity can be mediated by Na_V_ channels or by voltage-gated calcium channels, meaning TTX may or may not block their spiking (72, 73). Similar interchangeability is evident for the burst firing of Purkinje neurons (74).

Degeneracy also exists at the circuit level (75, 76), where it allows differences in the intrinsic excitability of component neurons to be offset (and effectively hidden) by differences in synaptic weights (77). Relevant for pain processing, the spinal dorsal horn circuit can achieve similar output using different synaptic weight combinations (78); specific neuron types may have a greater or lesser impact on circuit function depending on those weights. In effect, degeneracy introduces contingencies. The role of any ion channel in a neuron (or any neuron in a circuit) depends on the other ion channels in that neuron (or the synaptic connections with other neurons in the circuit). Because of such contingencies, a drug may engage its target without producing the intended cellular, circuit or clinical effect. Indeed, different combinations of GABA_A_ receptor activation and chloride driving force can produce equivalent synaptic inhibition (79), but when inhibition is incompensably compromised, the underlying cause necessitates different interventions (80). By this logic, if upregulation of Na_V_1.7 is responsible for nociceptor hyperexcitability after nerve injury or inflammation, Na_V_1.7 is an ideal target since “normal” neurons (not reliant on Na_V_1.7) would be spared the effects of a Na_V_1.7-selective drug, but the long-term efficacy of such a drug hinges on hyperexcitability remaining Na_V_1.7-dependent, which cannot be assumed (10). Furthermore, if myelinated afferents (which express minimal Na_V_1.7 (81)) are responsible for mechanical allodynia under neuropathic conditions (82-85), then Na_V_1.7-selective drugs should not be expected to alleviate that symptom, which evidently they do not (26), at least not through a direct mechanism. Indeed, ablating nociceptors abolishes acute and inflammatory pain but not neuropathic pain (23, 86). Pathological pain being mediated by more than one afferent type is another example of circuit-level degeneracy.

To be interchangeable, Na_V_ subtypes must functionally overlap (87, 88). Indeed, Na_V_1.8 and Na_V_1.7 are similar but not identical in their gating properties; for example, their voltage-dependencies partially overlap but the activation curve for Na_V_1.8 is right-shifted compared to Na_V_1.7 (89). Consequently, Na_V_1.7 activates at voltages near threshold whereas Na_V_1.8 tends to activate at suprathreshold voltages, during initiation and upstroke of the spike, respectively (34, 56). But that separation is not absolute. We found that Na_V_1.7 contributes to initiation of the first spike in DIV0 neurons, but because it inactivates more readily than Na_V_1.8, initiation of all subsequent spikes depends on Na_V_1.8 (see Fig. 5), which activates at perithreshold voltages because voltage threshold is high (depolarized) in the absence of Na_V_1.7. At DIV4-7, Na_V_1.7 still inactivates (which causes voltage threshold to rise) but, because of its higher density, continues to produce enough inward current to continue to initiate later spikes. The activation pattern we report for the first spike at DIV0 is consistent with Blair and Bean (90) who quantified the contribution of different Na_V_ channels by recording pharmacologically isolated currents while varying the holding potential according to the spike waveform. Our results go further in showing how responsibilities shift across different spikes within a train (because of differential Na_V_ inactivation) and across conditions (because of changes in Na_V_ expression). Although we focused on Na_V_ channels in this study, other ion channels are likely also undergoing changes; indeed, changes in the AHP shape between DIV0 and DIV 4-7 (see Figure 1A) point to changes in potassium channels. All neurons presumably regulate their excitability in a degenerate manner but likely do so by adjusting different sets of ion channels. Even within a given neuron type, the identify of adjusted ion channels may depend on the nature of the perturbation or on a multitude of other conditions.

With respect to reproducibility, labs testing nociceptors after different times in vitro would be expected to reach contradictory conclusions about the relative importance of a given Na_V_ subtype. Likewise, a testing protocol focusing on single spikes (the equivalent to the first spike in a train) would yield different results from one that considers repetitive spiking. Along the same lines, voltage clamp protocols that deliberately hold membrane potential at unnaturally hyperpolarized voltages to relieve inactivation before stepping up the voltage can give a misleading impression of how much a Na_V_ subtypes contributes under natural conditions (i.e. with natural levels of inactivation). Such discrepancies might be chalked up to irreproducibility if the consequences of those methodological differences are not appreciated, especially if one overlooks how degeneracy allows responsibilities to shift between ion channels. Indeed, the pain literature is replete with apparent inconsistencies. We would argue that most of those studies are correct, but only under limited conditions. Failure to identify and report those conditions (contingencies) represents a huge impediment to translation. A recent review of degeneracy in epilepsy (91) reveals many similarities with chronic pain. Observations that effective antiseizure medications often act on multiple targets and that patients with the same type of epilepsy respond heterogeneosly to a given treatment offer circumstantial evidence of degeneracy, but more deliberate testing is required to quantify degeneracy and its impact.

In summary, our results show that nociceptors can achieve equivalent excitability using different Na_V_ subtypes. The importance of a given subtype can shift on long and short timescales, yielding results that are seemingly inconsistent. By elucidating those shifting responsibilities, our results highlight the degenerate nature of nociceptor excitability and its functional implications. Degeneracy makes it impossible to claim without reservation that a particular Na_V_ subtype is uniquely responsible for pathological pain. Greater appreciation of degeneracy’s implications would prompt better experimental design, more cautious interpretation, and, ultimately, improved translation.

## MATERIALS AND METHODS

### Animals

All animal procedures were approved by the Animal Care Committee at The Hospital for Sick Children (protocol #53451) and were conducted in accordance with guidelines from the Canadian Council on Animal Care. We used the Cre-*lox*P recombinase system to generate mice that express ChR2-eYFP in Na_V_1.8-expressing neurons. Mice were obtained by crossing homozygous Ai32 mice (B6.Cg-Gt(ROSA)26Sortm32(CAG-COP4*H134R/EYFP)Hze/J) from Jax (#012569), which express ChR2(H134R)-eYFP in the presence of Cre recombinase, with Na_V_1.8-Cre mice (Tg(Scn10a-cre)1Rkun), which express Cre recombinase in Na_v_1.8-expressing neurons (kindly provided by Rohini Kuner). These neurons are primarily nociceptive and thermoreceptive (92). The Na_V_1.8 promoter leads to transgene expression in >90% of neurons expressing markers of nociceptors (21). To ensure that our transgenic mice were typical of wild-type mice with the same background (C57BL/6j), experiments reported in Fig. 1 were repeated in both genotypes for comparison. There was no effect of genotype on rheobase, spike height, input resistance, or spiking pattern, nor was there any significant interaction between genotype and effects of TTX except for spike height at DIV4-7, where TTX had a marginally larger effect in wild-type mice (Two-way ANOVA, F_1,54_=4.968, p=0.03, see source data file); therefore, we pooled the data for Fig.1. Having verified that our foundational observations held across different genotypes, we used transgenic mice for all subsequent experiments in order to identify eYFP-expressing nociceptors for patching, collection, or imaging.

### Dorsal root ganglia neuron cultures

All key reagents are listed in **Supplementary Table 2**. Methods for primary DRG culture have been described previously (93). Briefly, adult mice (> 7 week-old) were anaesthetised with isoflurane and perfused intracardiacally with cold HBBS (without Ca and Mg, LifeTech 14170112) supplemented with (in mM) 15 HEPES, 28 Glucose, 111 sucrose, and pH adjusted with NaOH to 7.3-7.4; osmolarity 319-321. Lumbar dorsal root ganglia (DRGs) were extracted (L2-5, except for CFA-inflamed mice, in which we only took L4), digested with papain (Worthington Biochemical Corp.) and collagenase (Worthington Biochemical Corp.)/dispase II (Sigma), and mechanically dissociated by trituration before being plated onto poly-D lysine-coated coverslips and incubated in Neurobasal™ media (Gibco 21103-049) supplemented with 1 % fetal bovine serum (FBS), B-27™ supplement (Thermo Fisher 17504-044) and 0.5mM L-Glutamine (Gibco 25030-081) for an initial period of 2 hours. After this, media was changed to maintenance media (same as plating media but without FBS) and cells were maintained in a 5% CO_2_ incubator at 37°C. Media was changed every 3-4 days thereafter. Neurons were recorded at two time points after plating: 2-8 hours (referred to as DIV0) or 4-7 days (referred to as DIV4-7). Neurons were tested at intermediate time points (DIV1-3) in the exploratory phase of our study but cellular heterogeneity prevented a clear picture of their TTX-sensitivity, presumably because different neurons shift from Na_V_1.8 to Na_V_1.7 at different rates. By DIV4, TTX sensitivity had stabilized, and so recordings from DIV4-7 were pooled for comparison with recordings on DIV0. When tests were conducted on a specific day within the DIV4-7 range, the specific DIV is reported but should be considered representative of the DIV4-7 range. Additional experiments are required to determine the exact time course of the Na_V_ switch and its mechanistic basis; to mitigate effects of cellular heterogeneity, this would ideally involve techniques that allow longitudinal measurements in the same cell.

### Electrophysiology

Coverslips with cultured neurons were transferred from the incubator to a recording chamber perfused with artificial cerebrospinal fluid containing (in mM): 126 NaCl, 2.5 KCl, 2.0 CaCl_2_, 1.25 NaH_2_PO_4_, 26 NaHCO_3_, 2 MgCl_2,_ and 10 glucose, bubbled with carbogen (5% CO_2_:95% O_2_) at room temperature. Neurons were visualized with gradient contrast optics on a Zeiss AxioExaminer microscope using a 40x, 0.75 NA water immersion objective (N-Achroplan, Zeiss) and IR-1000 Infrared CCD camera (Dage-MTI). YFP expression was visualized by epifluorescence (X-Cite, Excelitas) using a Zeiss filter set (46HE). A long-pass filter (OG590) was positioned in the transmitted light path to avoid activating ChR2 while patching. No optogenetic testing was performed as part of this study. Cells expressing YFP and with a soma diameter <25 µm were targeted for whole cell recording using pipettes (∼5 MΩ resistance) pulled from borosilicate glass (WPI).

For current clamp recordings, pipettes were filled with intracellular solution containing (in Mm): 140 K-gluconate, 2 MgCl_2_, 10 HEPES, 0.2 EGTA, 3.8 Na-ATP and 0.4 Na-GTP with pH adjusted to 7.3 with KOH; osmolarity was ∼300 mOsm. A liquid junction potential correction of 15 mV was applied to all reported voltages. Series resistance was compensated to >70%. Signals were amplified with an Axopatch 200B amplifier (Molecular Devices, Sunnyvale, USA), low-pass filtered at 2 kHz, digitized with a Power1401 A/D device (Cambridge Electric Design, Cambridge, UK), and recorded at 10 kHz using CED software Signal version 6. After the natural resting membrane potential was noted, neurons were adjusted to -70 mV using continuous current injection in current clamp mode. Action potentials (spikes) were evoked using a series of 1-second long depolarizing current injections. Rheobase was defined as the minimal current required to evoke a spike. Neurons were tested with current injections from 1x rheobase to 4x rheobase using increments of 0.5x rheobase. Repetitive spiking neurons were defined as those producing ≥3 spikes in response to any stimulus intensity; transient spiking neurons consistently produced ≤2 spikes. Spike threshold was defined as voltage where dV/dt first exceeds 5 mV/ms (94). Spike height was measured from threshold to peak of the action potential. Only neurons with a resting membrane potential below -45 mV, spikes overshooting 0 mV and recordings with <20% change in series resistance were tested and analyzed. For dynamic clamp experiments, the pipette shank was painted with Sylgard (Dow) to reduce pipette capacitance. Virtual Na_V_1.7 and Na_V_1.8 conductances were introduced into the cells using CED software Signal v6. Currents were defined using the Hodgkin-Huxley equation, using the same parameter values as in our computational model (see below).

For voltage-clamp recordings, the bath solution was adjusted to reduce sodium currents to ensure proper clamping (85). Bath solution contained (in mM): 65 NaCl, 50 choline chloride, 5 KCl, 5 HEPES, 5 MgCl_2_, 10 glucose, and 0.1 CaCl_2_, plus 0.1 CdCl_2_ to block calcium currents, and 20 TEA and 5 4-AP to block potassium currents; pH was adjusted to 7.4 with NaOH. Pipettes were filled with intracellular solution containing (in Mm): 140 CsCl, 10 HEPES, 2 MgCl_2_, 1 EGTA, 3.8 Na-ATP, 0.4 Na-GTP; pH was adjusted to 7.3 with CsOH. The resulting pipette resistance was ∼3 MΩ. A liquid junction potential correction of 4.8 mV was applied to all command voltages. Sodium currents were recorded during 20 ms-long steps from -85 mV to voltages between -45 and +15 mV. Series resistance was compensated to >80%. Signals were amplified, low-pass filtered at 5 kHz, and digitized as described for current clamp recordings.

For all in vitro pharmacology, drugs were bath applied at a concentration chosen to selectively block the Na_V_ subtype of interest based on published EC50 values (see Supplementary Table 3)

### Quantitative reverse transcription PCR (RT-qPCR)

Cultured DRG neurons <25 μm and expressing eYFP were identified as described above for patching. Coverslips were perfused with aCSF made with DEPC-treated ddH_2_O, and identified neurons were collected using a glass pipette filled with intracellular solution also made from DEPC-treated ddH_2_O (composition otherwise the same as described above for electrophysiology). Approximately 50 neurons were collected at DIV0 and at DIV4-7. Total mRNA was extracted with a PureLink RNA mini kit after digestion of genomic DNA with DNase I (Thermo Fisher Scientific) and the cDNA was synthesized with a SuperScript II first-strand synthesis kit (Thermo Fisher Scientific) according to instructions. RT-qPCR was performed with the cDNA primers of target genes (**Supplementary Table 3**), and the PowerUp SYBR^®^ Green master mix (Thermo Fisher Scientific) in the QuantStudio-3 real-time PCR system. The primers were designed with IDT and spanned at least one exon longer than 1000 bp in order to exclude contamination from genomic DNA. Non-RT mRNA was also used as a negative control to exclude contamination from genomic DNA. All target genes were performed in triplicate for each sample and the experiments were repeated at least 3 times. Na_V_1.7 and Na_V_1.8 transcript levels were analyzed using the 2-ΔΔCT method and compared with the housekeeping gene HPRT.

### Immunocytochemistry

Cultured DRG neurons were treated with 4% paraformaldehyde for 10 minutes, rinsed 3x with cold PBS, and permeabilized with 0.1% Triton X-100 in PBS. After another 3x rinse with PBS, neurons were treated with 10% normal goat serum for 30 min followed with rabbit primary Na_V_1.7 antibody (1:200, ASC-008, Alomone) or Na_V_1.8 antibody (1:200, ASC-028, Alomone) in PBS with 0.1% Tritween-20 and 1% BSA for 1 h. For some of the coverslips, primary antibodies were replaced with control peptides (ASC008AG1040 for Na_V_1.7 and ASC016AG0640 for Na_V_1.8) provided by Alomone as negative controls. Following 3x rinse in PBS, neurons were incubated in the dark with goat anti-rabbit secondary antibody Alexa Fluor-647 (1:500, Abcam) in PBS containing 1% BSA for 1 h, followed by DAPI staining for 10 min. All incubations were done at room temperature. Finally, coverslips were mounted on slides with mounting media (Abcam, ab128982), imaged with a spinning disk confocal microscope (Quorum Technologies) using the same acquisition setting across all imaging sessions, and analyzed with Volocity software (v6.5.1). Protein levels are measured using fluorescence intensity and expressed relative to each other (e.g. ratios in Fig 6C) or relative to fluorescence intensity for YFP in the same cells. Each condition was tested in a minimum of 3 animals.

### Behavioral testing

Behavioral tests were performed on adult mice (male and female, 8-12 weeks). Mice were acclimated to the testing environment for at least 1 h the day prior to start of experiments. Behavioral testing (von Frey test and Hargreaves test) was then performed for 2-3 consecutive days for baseline and for another 3 days after CFA injection. Behavioral tests were performed at the same time in the morning, at room temperature (21°C) following a one-hour acclimation period. Animals were randomly assigned to experimental groups and the experimenter was blind to the drug condition.

*CFA injection*. CFA (Sigma, F5881) was thoroughly dissolved in saline (1:1) by vortexing the mixture. The resulting CFA solution (20 µl) was injected subdermally into the left hind paw under light isoflurane anaesthesia. The injection was performed shortly after the last baseline test, on Day 0.

*PF-71 administration*. Injectable PF-71 solution was prepared by first dissolving PF-71 in DMSO to make a 5% stock solution; dissolution was achieved by heating to 37°C and vortexing. On the day of injection, stock solution was dissolved in sunflower oil (5% v/v) by sonicating for 5 min. Freshly prepared final PF-71 solution was injected intraperitoneally (1 g/kg body weight). Behavioral testing was performed 2 hours after injection of PF-71 or vehicle.

*Von Frey testing*. Mechanical hyperalgesia was assessed with von Frey filaments (North Coast) using the SUDO method (95). The average of 3 trials in each animal was used for analysis.

*Hargreaves testing.* Thermal hyperalgesia was assessed with the Hargreaves apparatus (Ugo Basile, Italy). Radiant heat was applied to the plantar surface of the left hind paw. Interval between stimulus onset and paw withdrawal was defined as paw withdrawal latency (PWL). A 20 s cut-off prevented damage to the skin if the animal failed to withdraw. The average of 3 trials in each animal was used for analysis.

### Statistical analysis

Results are expressed as mean ± SEM when data are normally distributed or otherwise as median and quartiles. Normality was tested using the Kolmogorov-Smirnov test. Analysis was performed with GraphPad Prism (v9) and SigmaPlot (v11). Normally distributed data were compared using t-tests or two-way ANOVAs followed by a Student Newman-Keuls post hoc test. Non-normally distributed data were compared using Mann-Whitney and Wilcoxon signed rank tests. Fisher exact and McNemar test were used for categorical data. Exact significance values and test results are reported throughout figure legends.

### Computer model

Two separate, single compartmental models were built for DIV 0 and DIV4-7. They have the same, seven conductances: *g*_Nav1.3_, *g*_Nav1.7_, *g*_Nav1.8_, *g*_Kdr_, *g*_M_, *g*_AHP_ and *g*_Leak_. Channel equations and their gating parameters are provided in **Supplementary Table 4**. Conductance densities at baseline (**Supplementary Table 5**) were tuned to qualitatively reproduce the changes in Na_V_ channel expressions at DIV 0 and 4-7 indicated by the experiments; changes to other channels were minimized between the two models. The effects of ICA, PF-71 and PF-24 were simulated by adjusting *ḡ*_Nav1.3_, *ḡ*_Nav1.7_ and *ḡ*_Nav1.8_, respectively, as reported in the figures. To re-introduce voltage noise that is otherwise missing from simulations, we included an Ornstein-Uhlenbeck process with *μ*_noise_ = 0 µA/cm^2^, *σ*_noise_ = 0.05 µA/cm^2^, and *τ* = 5 ms. All computer code is available at ModelDB (http://modeldb.yale.edu/267560; password: excitability). All simulations were conducted in MATLAB using the forward Euler integration method and a time step of 0.05–0.1 ms.

## CONTRIBUTIONS

YX, SR, SAP designed the research; YX, JY collected data, YX, JY, SR, SAP analyzed data; YX, SR, SAP wrote the manuscript.

## ACKNOWLEDGMENTS

This work was supported by a Restracomp fellowship to JY and by a Canadian Institutes of Health Research Foundation Grant (FDN167276) to SAP. We thank Rohini Kuner for providing Na_v_1.8-Cre mice, Jason Jeong and Russell Smith for expert technical assistance with animal care and cell cultures, and Yongqian Wang for advice on qPCR data acquisition.

---

**Figure 2 – figure supplement 1.**
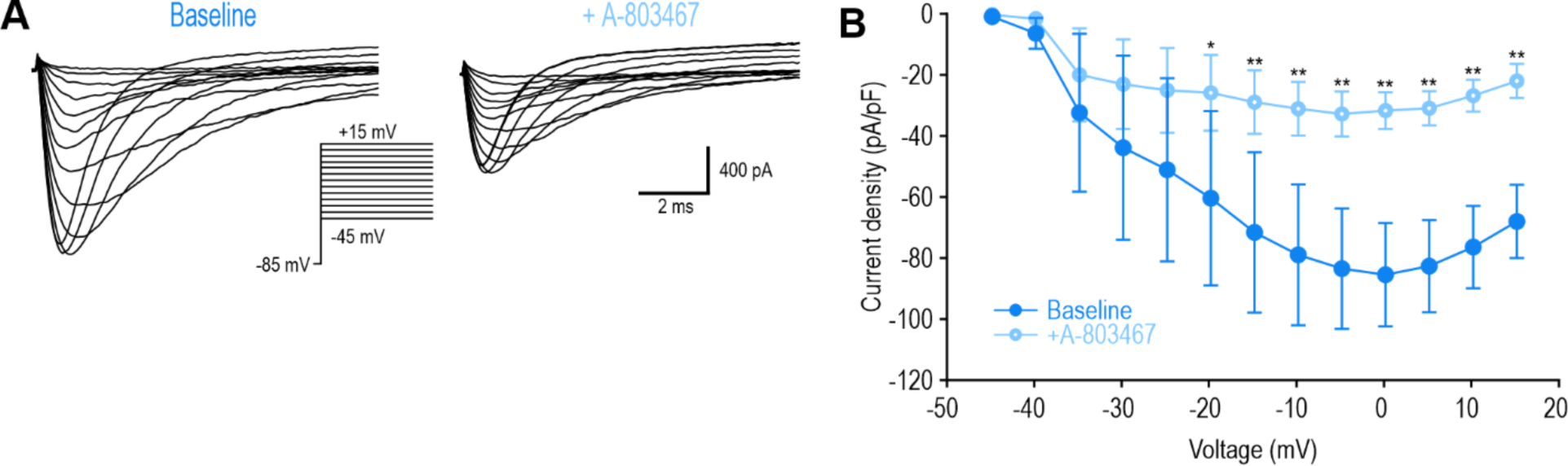
Inhibiting Na_V_1.8 at DIV0 with A-803467 had the same effect as PF-24. **(A)** Sample voltage clamp recording at DIV0 before and after A-803467 (1 µM). **(B)** Peak current was significantly reduced by A-803467 (F_1,84_=9.935, p=0.016, two-way RM ANOVA, n=8). *, p<0.05; **, p<0.01; Student-Newman-Keuls post-hoc test in B.

**Figure 2 – figure supplement 2.**
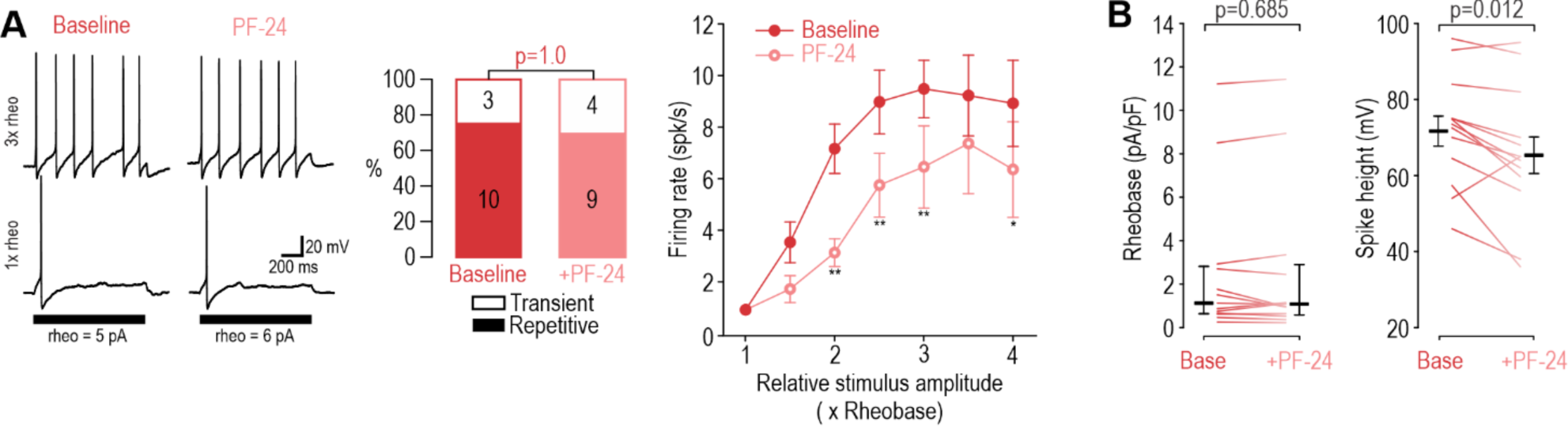
Inhibiting Na_V_1.8 with PF-24 at DIV4-7 had negligible effects. **(A)** Inhibiting Na_v_1.8 with PF-24 (1 µM) did not affect spiking pattern (χ^2^ =0.00, p=1.00, McNemar test) and modestly reduced firing rate (F_1,54_=9.745, p= 0.012, two-way RM ANOVA, n=10) in DIV4-7 neurons. **(B)** PF-24 did not affect rheobase (Z_12_=0.420, p=0.685, Wilcoxon Rank test) but did reduce spike height (T12=2.939, p=0.012, paired-t-test). *, p<0.05; **, p<0.01; Student-Newman-Keuls post-hoc tests in A.

**Figure 3 – figure supplement 1.**
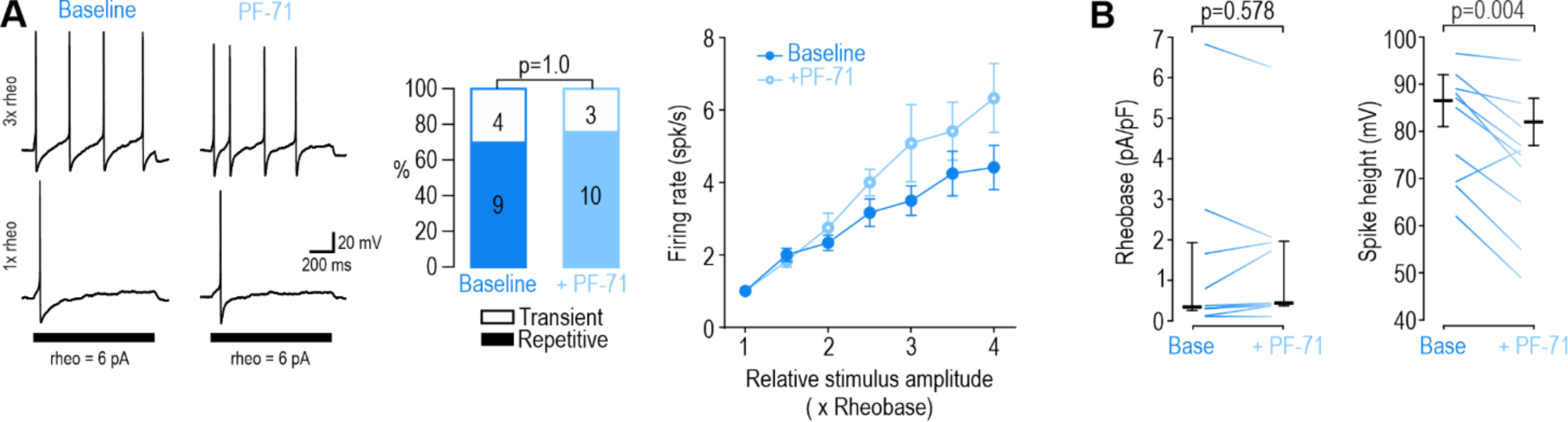
Inhibiting Na_V_1.7 at DIV0 had negligible effects. **(A)** Inhibiting Na_V_1.7 with PF-71 (30 nM) did not alter spiking pattern (χ^2^=0.00, p=1.00, McNemar test) or reduce firing rate (F_1,30_=5.805, p=0.061, two-way RM ANOVA, n=6) in DIV0 neurons; in fact, firing rate was slightly increased. **(D-E)** PF-71 did not affect rheobase (Z_9_=0.677, p=0.578, Wilcoxon rank test) but did reduce spike height (T_9_=3.759, p=0.004, paired-t-test).

**Figure 3 – figure supplement 2.**
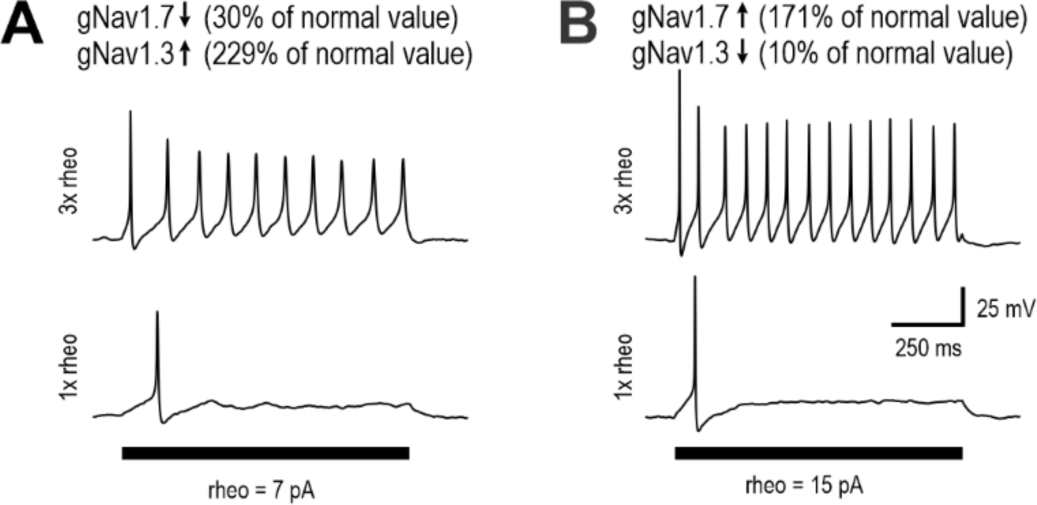
Na_V_1.7 and Na_V_1.3 currents can compensate for each other. **(A)** In the DIV4-7 model, reducing *g*_Nav1.7_ to 30% of its normal value (35→10 mS/cm^2^) can be compensated for by increasing *g*_Nav1.3_ to 229% of its normal value (0.35→0.8 mS/cm^2^) to maintain repetitive spiking. **(B)** Conversely, reducing *g*_Nav1.3_ to 10% of its normal value (0.35→0.035 mS/cm^2^) can be compensated for by increasing *g*_Nav1.7_ to 171% of its normal value (35→60 mS/cm^2^) and maintain repetitive spiking.

**Figure 4 – figure supplement 1.**
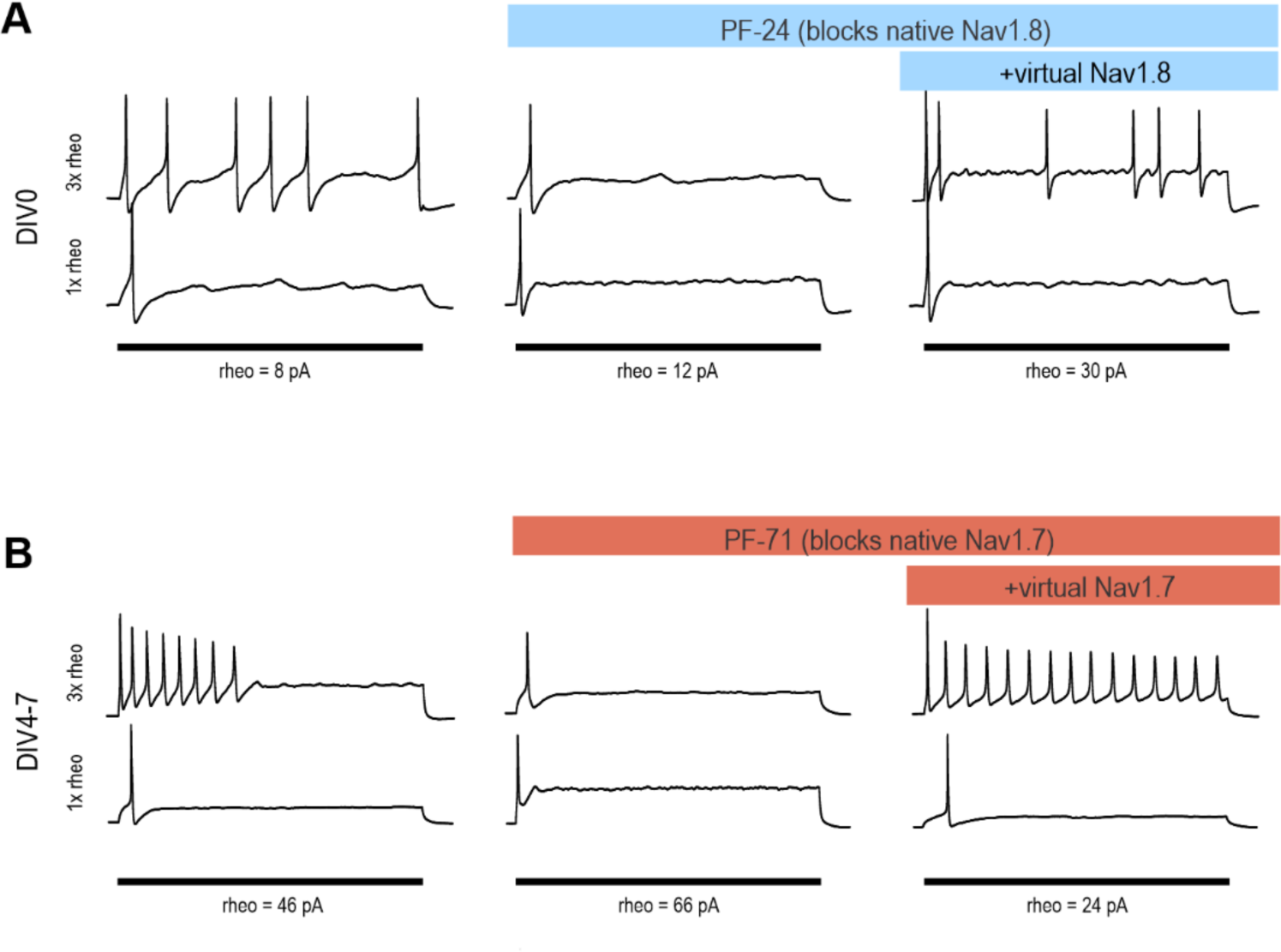
Virtual conductances restored repetitive spiking after pharmacological inhibition of the corresponding native conductance had converted the neuron to transient spiking. **(A)** Sample response at DIV0 showing that a virtual Na_V_1.8 conductance applied with dynamic clamp restored repetitive spiking after inhibiting native Na_V_1.8 channels with PF-24. This restoration was repeated in 3 of 3 neurons tested. **(B)** Sample recording at DIV4-7 showing that a virtual Na_V_1.7 conductance restored repetitive spiking after inhibiting native Na_V_1.7 channels with PF-71. This restoration was repeated in 4 of 4 neurons tested.

**Figure 5 – figure supplement 1.**
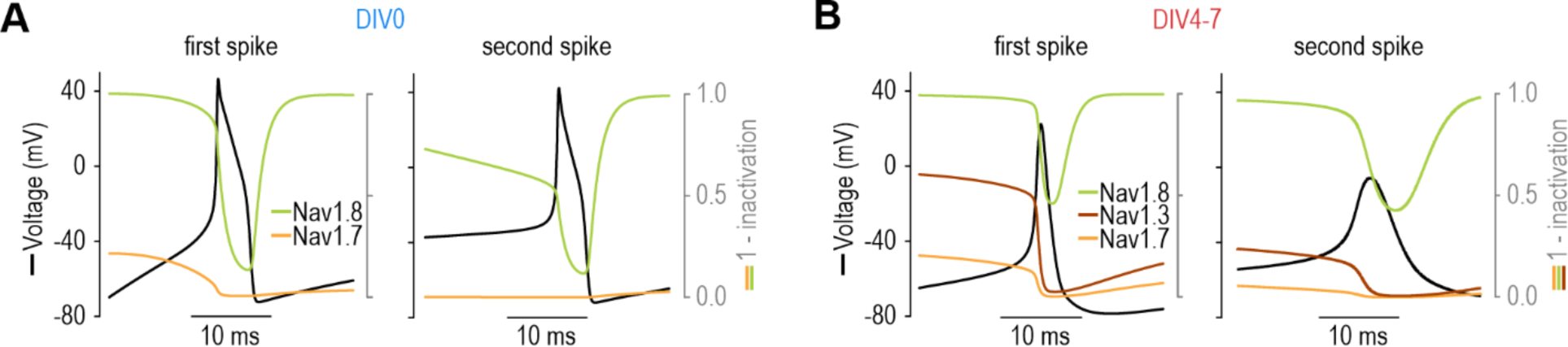
Channel inactivation affects Na_V_ subtype contribution on short timescale. **(A)** In the DIV0 model, Na_V_1.7 contributed to the first spike but its inactivation meant that all subsequent spikes relied exclusively on Na_V_1.8. **(B)** In the DIV4-7 model, despite some inactivation of Na_V_1.3 (maroon) and Na_V_1.7 (orange), the remaining current was still large enough (because of the higher *g*_max_ of those two subtypes) to produce inward current sufficient to support repetitive spiking despite the low *g*_max_ of Na_V_1.8 in the DIV4-7 model.

**Figure 5 – figure supplement 2.**
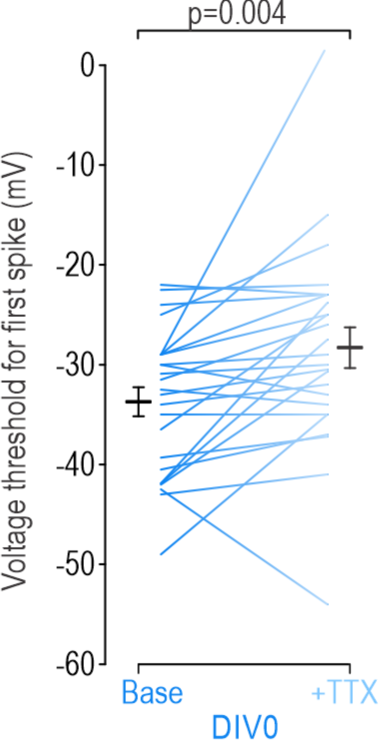
Effect of TTX on voltage threshold in DIV0 neurons. Despite TTX having negligible effects in DIV0 neurons according to our initial analysis (see Fig. 1), simulation results in Fig. 5A,B predicted that the first spike was nonetheless initiated by Na_V_1.7. By extension, this predicted that TTX should cause a depolarizing shift in voltage threshold for the first spike. Analysis of the experimental data confirmed this to be true, with threshold (mean±SEM) increasing from -33.7±1.4 mV at baseline to -28.3±1.4 mV after TTX (T_24_=-3.19, p=0.004, paired t-test). Confirmation of this unexpected prediction helps further validate our model neuron.

**Supplementary Table 1.** Model data before and after channel “inhibition”.

|  | **DIV0** |  | **DIV4-7** |  |  |
| --- | --- | --- | --- | --- | --- |
|  | Baseline | ▼ Na<sub>v</sub>1.8 | Baseline | ▼ Na<sub>v</sub>1.7 | ▼ Na<sub>v</sub>1.3 |
| **Rheobase (pA)** | 17 | 22 | 12 | 16 | 21 |
| **Spike height (mV)** | 72.8 | 46.9 | 57.1 | 21.4 | 52 |
| **RMP (mV)** | -69.0 | -69.7 | -70.0 | -69.8 | -70.4 |

**Supplementary Table 2.** Reagents.

| Reagent | Description | Source |
| --- | --- | --- |
| **TTX** | Tetrodotoxin citrate | Alomone |
| **PF-24** | PF-01247324 | Sigma |
| **PF-71** | PF-05089771 | Alomone |
| **ICA** | ICA-121431 | Tocris |
| **Papain** | Papain lyophilized power | Worthington |
| **Collagenase** | Collagenase type II | Worthington |
| **Dispase II** | Dispase type II | Sigma |

**Supplementary Table 3.** EC50 values.

| Drug (concentration used) | EC50 values |  |  | References |
| --- | --- | --- | --- | --- |
|  | Na<sub>v</sub>1.3 | Na<sub>v</sub>1.7 | Na<sub>v</sub>1.8 |  |
| **TTX (100 nM)** | **4 nM** | **2 nM** | 46±10 μM | (21, 96) |
| **ICA-121431 (1 μM)** | **20 nM** | >10 μM | >10 μM | (53) |
| **PF05089771 (30 nM)** | 11 μM | **11 nM** | >10 μM | (40) |
| **PF-01247324 (1 μM)** | >30 μM | >30 μM | **331 nM** | (50) |
| **A-803467 (1 μM)** | 2.5 μM | 6.7 μM | **8 nM** | (97) |

Bolded entries indicate the Na<sub>v</sub> subtype that is selectively blocked

**Supplementary Table 4.**
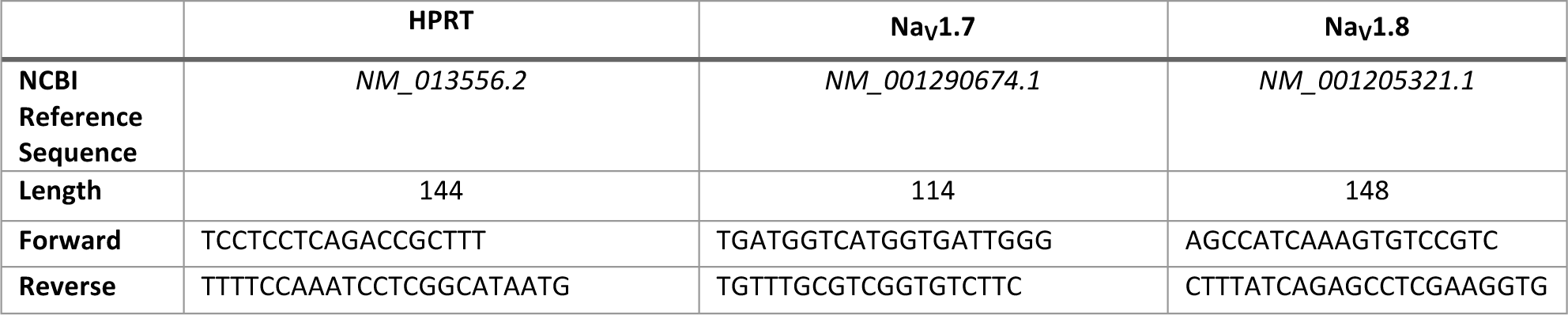
Primers.

**Supplementary Table 5.**
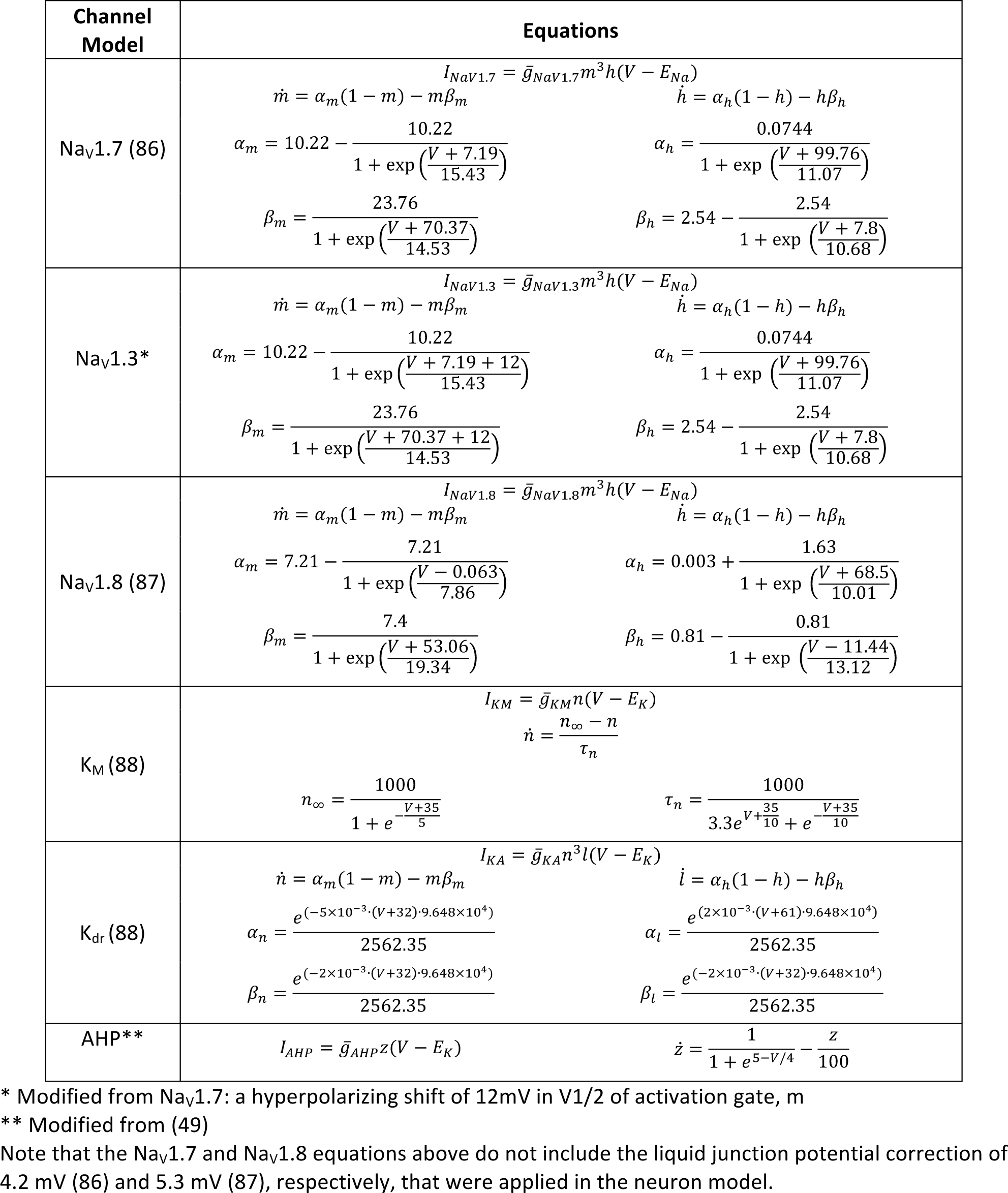
Model equations.

**Supplementary Table 6.** Conductance densities at baseline for DIV 0 and 4-7 models.

|  | Conductance densities at baseline (mS/cm<sup>2</sup>) |  |  |  |  |  |  |
| --- | --- | --- | --- | --- | --- | --- | --- |
| | $\bar{g}_{\text{Nav1.3}}$ | $\bar{g}_{\text{Nav1.7}}$ | $\bar{g}_{\text{Nav1.8}}$ | $\bar{g}_{\text{Kdr}}$ | $\bar{g}_{\text{M}}$ | $\bar{g}_{\text{AHP}}$ | $\bar{g}_{\text{Leak}}$ |
| DIV 0 | 0 | 3 | 30 | 3 | 0.05 | 3.5 | 0.025 |
| DIV 7 | 0.35 | 35 | 0.2 | 3.5 | 0.5 | 2.5 | 0.035 |

## Notes

### Competing Interest Statement

SAP has served on the Scientific Advisory Boards of Boston Scientific and Presidio Medical and has received grant funding from Boston Scientific and Presidio Medical.

### Summary of Updates

The manuscript has been revised in response to eLife reviewer comments.

